# Distinct roles for DNA- and RNA-sensing immune pathways in control of genital HSV-2 infection and restriction of neuroinvasion

**DOI:** 10.64898/2026.07.02.735977

**Authors:** Xin Lai, Xiangning Ding, Ryo Narita, Alexander Schmitz, Mathias Franzén Boger, Gudrun Winther, Marie B Iversen, Giorgia Marino, Sara R N Jensen, Kasper Thorsen, Ole A Ahlgreen, Christian Bjerggaard Vægter, Amanda Eskelund, Faezeh Darki, Thierry P.P. van den Bosch, Klara Hasselrot, Kristina Broliden, Line S Reinert, Søren R Paludan

## Abstract

Early control of viral infection is thought to rely on pattern-recognition receptors (PRRs) that induce interferons (IFNs) and leukocyte recruitment, but how distinct PRRs coordinate mucosal antiviral defense remains unclear. We show that both the DNA-sensing cGAS–STING pathway and the RNA-sensing RLR–MAVS pathway are required for protection upon genital herpes simplex virus type 2 (HSV-2) infection. cGAS deficiency increased infection-induced pathology in both epithelial and submucosal compartments, whereas MAVS deficiency primarily affected the epithelium. Spatial proteomics and regional transcriptomics revealed that cGAS was essential for early epithelial TBK1 activation, expression of IFN-stimulated genes and recruitment and activation of myeloid and lymphoid cells to the epithelium. While MAVS was essential for full TBK1 activation it had limited impact on the induced IFN response. However, MAVS sustained basal epithelial expression of the antiviral factors IFITM1 and 3, which exert antiviral activity against HSV-2. Notably, cGAS deficiency impaired submucosal IFN responses and enabled viral spread into this tissue, enabling infection of intervening neurons and dissemination to the central nervous system. These findings define coordinated, compartment-specific innate defense against infections.

## INTRODUCTION

Herpes simplex virus (HSV-2) is the main cause of genital herpes (*1, 2*), which is a sexually-transmitted disease leading to genital ulcers. Following infection with HSV-2 in epithelial cells (ECs) in the genitals, the virus establishes latent infection in dorsal root ganglia (DRG), enabling periodic reactivation with risk of reoccurrence of ulcers. Importantly, pregnant women with genital herpes are at risk of transmitting the infection to their child during delivery through the birth canal, leading to neonatal herpes, which is a very serious condition (*2*). Moreover, in infected individual, the virus can gain access to the central nervous system (CNS), through the neuronal route, and give rise to serious complications, including myelitis and meningitis (*3*). Therefore, host immune activities against genital HSV-2 infection needs to control the infection, spare healthy tissue, and limit viral access to neuronal tissue.

Nucleic acid sensing is important for early immune control of viruses, including HSV infections. Mammalian cells encode pattern recognition receptors (PRRs) detecting RNA and DNA (*4, 5*). For instance, among the Toll like receptors (TLRs), which localizes to plasma or endosomal membranes, TLR3 senses double-stranded (ds) RNA and TLR9 senses DNA (*6, 7*). In addition, a number of cytoplasmic nucleic acid sensors also exist. This includes the RNA-sensing RIG-like receptors (RLR), which signal through the adaptor protein mitochondrial antiviral-signaling protein (MAVS) (*8, 9*), and the DNA-sensing cyclic GMP-AMP synthase (cGAS) pathway, which signals through the adaptor protein Stimulator of interferon (IFN) genes (STING) (*10, 11*). A central downstream activity induced by PRRs is the expression and secretion of type I and III interferons (IFNs), which are cytokines with potent antiviral activity. In addition, PRR activation induces expression of chemokines and recruitment of leukocyte populations to the site of infection. These cells are essential for control of infection but can also contribute to amplification of pathology. For instance, natural killer (NK) cells are recruited to the vagina following HSV-2 infection in mice (*12*), and have been reported to be involved in control of infection (*13*). Conversely, recruitment of macrophage can amplify inflammation (*14*), and potentially also predispose to co-infection with human immunodeficiency virus (*15*).

While the majority of research on nucleic acid sensors in relation to control of virus infections has focused on sensors detecting the same form of nucleic acid as the genome of the virus under investigation (e.g. DNA sensors in the case of DNA viruses), it is becoming increasingly apparent that multiple pathways are activated in parallel during infections in tissues. For instance, dsRNA accumulates in permissive cell lines upon infection with several DNA viruses (*16*). Moreover, internal folding of viral or host RNAs have been reported to have immunostimulatory activity (*17–19*). Finally, mitochondrial dysfunction occurs during many DNA viral infections (*20*), and can lead to release of mitochondrial dsRNA species (*21, 22*). Consequently, DNA viruses have been reported to activate RNA sensors, TLR3 and RLRs (*17, 18, 23–25*). Likewise, infections with a range of RNA viruses are well-described to disturb mitochondrial function leading to release of DNA into the cytoplasm and activation of the cGAS-STING pathway (*26–29*). However, the physiological importance parallel activation of multiple PRRs during natural infections is not well explored.

Regarding the current understanding of nucleic acid-sensing PRRs in control of HSV-1 and 2, cGAS-STING is important for control of HSV-1 infection both on mucosal surfaces and in the CNS in mice (*11, 30–32*). In addition, lack of the TLR3 adaptor protein TRIF, and to a lesser extent also MAVS, have been reported to lead to elevated HSV-1 load in the CNS (*33*). In humans, deficiency in TLR3 and pathway-specific genes have been reported to be associated with herpes simplex encephalitis (HSE) (*34, 35*). Moreover, defects in genes encoding proteins shared between numerous PRRs and involved in the induction and function of IFNs predispose to HSE (*36–39*). Regarding HSV-2, cGAS has been reported to be important for control of genital infection in mice (*40*), which is highly dependent on the type I IFN system (*41*). We also reported a role for TLR3 in control of HSV-2 in the spinal cord in mice on a mixed B6-129S genetic background (*42*). In this system, astrocytes were the main cell type mediating TLR3-driven IFN responses. However, it remains unknown whether a broader panel of pathogen-sensing pathways are involved in sensing of HSV-2 in thein the female genital tract (FGT), and how they may mechanistically orchestrate the antiviral response to avoid immunopathology and limit viral spread.

In this study we report that cGAS and MAVS are both essential for full control of HSV-2 infection in the FGT in mice, thus demonstrating a role for both DNA and RNA sensing pathways in full innate protection. Both pathways are involved in control of HSV-2 in vaginal ECs, while only cGAS contributes to restriction of viral spread beyond the epithelium and ultimately neuroinvasion. Using spatial proteomics and transcriptomics technologies we show that cGAS drives the early induced IFN response in ECs, which extends into the stroma, and also enables the recruitment and activation of immune cells. By contrast, MAVS controls basal expression of a subsets of ISGs in ECs, thus imposing an immediate barrier for control of HSV replication and limits the activation of cGAS-STING-driven immune activation.

## RESULTS

### Host defense against genital HSV-2 relies on both RNA and DNA sensing pathways

In order to examine what innate immunological pathways that govern defense against genital HSV-2 infection and prevention of disease, we infected WT, *cGas^-/-^*, *Tlr3^-/-^*, and *Mavs^-/-^*mice with HSV-2 in the FGT (Fig. 1A). The sensors and adaptors for these pathways were expressed in the mouse FGT (Fig. S1A). The lack of cGAS rendered the mice highly susceptible to HSV-2 infection (Fig. 1B-C) whereas TLR3 was not essential for resistance at this infection dose (Fig. 1D-E). Interestingly, however, mice devoid of RLR signaling due to MAVS-deficiency also exhibited significantly elevated susceptibility to HSV-2 infection in the FGT (Fig. 1F-G). The effect of cGAS- and MAVS-deficiency was also observed at ten times higher infection doses, where TLR3-deficiency also exhibited minor impact on disease outcome (Fig. S1B-D), in agreement with previous results (*42*). When testing other models for HSV-1 and 2, we observed that MAVS-deficiency also significantly decreased survival after ocular HSV-1 or 2 infections (Fig. S1E-F).

**Figure 1.**
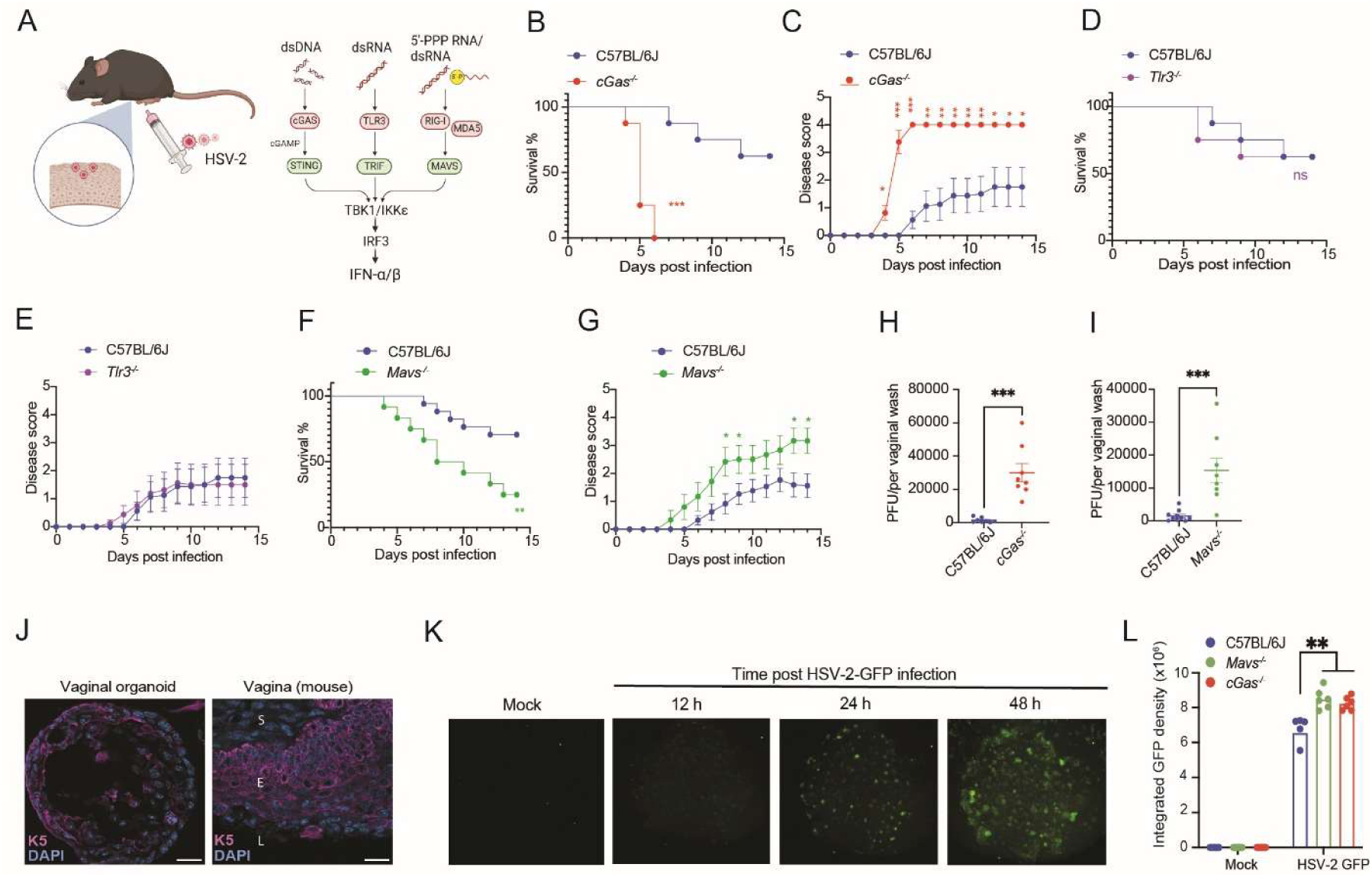
Host defense against genital HSV-2 relies on both RNA and DNA sensing pathways. (A) Illustration of mouse infection model, and immune pathways examined in the present study. The image was generated in Biorender. (B-G) Mice were infected intravaginally with 6.7 × 10^3^ plaque-forming units (p.f.u.) of HSV-2 (strain 333) and monitored on subsequent days for survival and disease scores. (B-C) WT (C57BL/6J) and *cGas^-/-^*(n = 8 mice/group), (D-E) WT and *Tlr3^-/-^* (n = 8 mice/group), and (F-G) WT and *Mavs^-/-^* (n = 4-8 mice/group). (H-I) WT, *cGas^-/-^*, and *Mavs^-/-^* mice were infected intravaginally with 6.7 × 10^4^ p.f.u. of HSV-2, and vaginal washes were collected on day 2 post infection for plaque assay (n = 8-10 mice/group). (J) Immunohistological staining of vaginal organoid and mouse vaginal tissue sections with epithelial cell marker cytokeratin 5 (K5, red) and DAPI (blue). Scale bar: 20 μm. (K-L) Visualization of mouse vaginal organoids infected with HSV-2 GFP at the indicated time points post infection, and quantification of the 24 h time point in WT, *cGas^-/-^*, and *Mavs^-/-^* organoids. (n=4-6 wells/ group)(B, D, F) Survival was analyzed using log-rank Mantel–Cox test. (C, E, G) Disease progression was analyzed using 2-way repeated-measures ANOVA with Sidak’s multiple-comparison test. (H and I) Wilcoxon rank-sum test. (L) 2-way repeated-measures ANOVA with Sidak’s multiple-comparison test *P* values were calculated using ANOVA with a Turkey’s post hoc test. Plots show means ± st.dev; ^∗^*p* < 0.05, ^∗∗^*p* < 0.01, ^∗∗∗^*p* < 0.001; ns, not significant.

To explore whether lack of cGAS or MAVS influenced HSV-2 replication in the FGT, we isolated vaginal washes from mice for plaque assay. Washes from *cGas^-/-^* and *Mavs^-/-^* mice contained significantly higher levels of HSV-2 compared to WT mice (Fig. 1H-I). To further examine the elevated susceptibility to HSV-2 infection in the vaginal epithelium, we generated mouse vagina organoids, which are composed of ECs (Fig. 1J, Fig S1G-H) and can be productively infected with HSV-2 (Fig. 1K). Importantly, the cGAS- and MAVS-deficient mouse vaginal organoids showed elevated viral gene expression compared to WT (Fig. 1L). Collectively, these data show that cGAS and MAVS play non-redundant roles in control of HSV-2 infection in the FGT and suggests they operate through epithelium-intrinsic vaginal antiviral mechanisms.

### Tissue pathology in the HSV-2-infected FGT

To examine the tissue distribution of HSV-2 in the FGT of the mouse strains under investigation, we stained vaginal sections for HSV-2. The immunohistochemical analysis of vaginal tissues revealed that the virus infection foci in the FGT of *cGas^-/-^* and *Mavs^-/-^* mice were larger and accompanied with more extensive disruption of the epithelial cell layer (Fig. 2A). However, the two gene-modified mouse strains differed. In the *Mavs^-/-^* mice, HSV-2 was limited to the epithelial layer, similarly to the WT mice, while there was wider spread of virus and into the deeper tissue in *cGas^-/-^* mice and tissue destruction (Fig. 2A).

**Figure 2.**
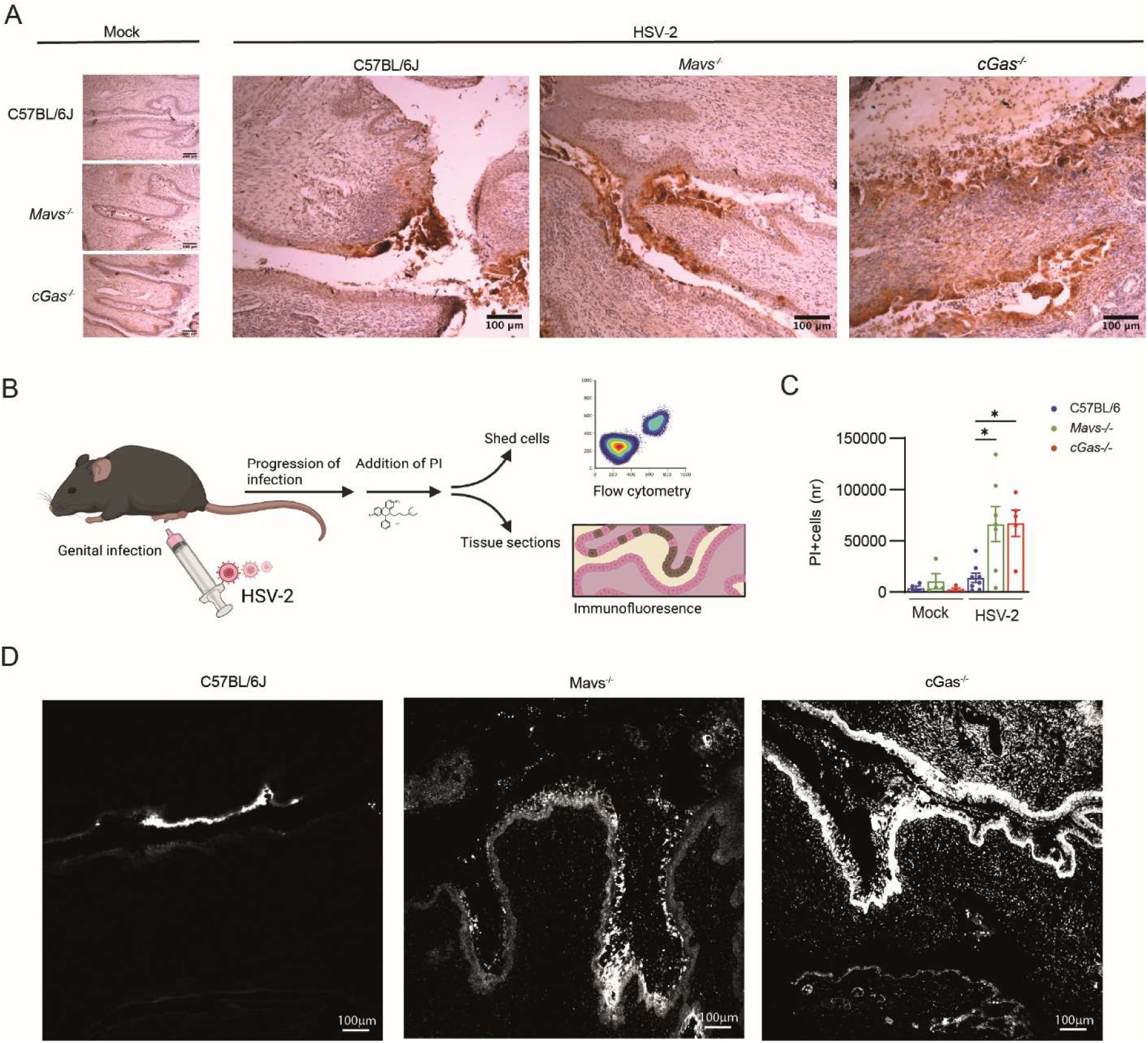
DNA and RNA-sensing pathways differentially affect tissue pathology in the mucosal area. (A) WT, *cGas^-/-^*, and *Mavs^-/-^*mice were infected intravaginally with 6.7 × 10^4^ p.f.u. of HSV-2 for 48 h. Tissue sections were immunostained with anti-HSV antibody. Scale bar: 100 μm. (n = 4-6 mice/group). (B) Graphical illustration of the experimental set-up to examine for cell death in the vaginal tissue. Made with Biorender. (C) Cells shed into the vaginal lumen were isolated 48 h p.i. after Propidium iodide (PI) treatment 10 min before vaginal wash. The nr of dead (PI^+^) cells among total Hoechst^+^ cells were quantified by flow cytometry. Numbers are shown as total number of dead PI^+^ cells detected per mouse (n = 4-8 mice/group). (D) Tissue-sections from mice infected for 48 h and treated with PI 10min before sacrifice were visualized by fluorescence microscopy. Scale bar: 100 μm. Images are shown in black and white. (n = 4-6 mice/group). (C) *P* values were calculated using ANOVA with a Turkey’s post hoc test. Plots show means ± st.dev; ^∗^*p* < 0.05.

To explore whether the distribution of cell death differed between the mouse strains, we treated mice intravaginally with the necrotic cell death marker propidium iodide (PI) before termination of experiments and isolated shedded cells from the vaginal lumen and visualized tissue sections (Fig. 2B). As expected, both *cGas^-/-^* and *Mavs^-/-^* mice showed elevated levels of shedding of necrotic cells into the vaginal lumen compared to WT mice after HSV-2 infection (Fig. 2C). This was paralleled by increased PI staining in the epithelial layer in both mouse strains (Fig. 2D). However, the profound spreading of cell death to the submucosal layer was observed only in *cGas^-/-^* mice (Fig. 2D). These data show that lack of either innate cytosolic RNA or DNA sensing pathways facilitates spread of HSV-2 in the vaginal epithelial layer, while only the cGAS-STING pathway is involved in limiting spread of virus and tissue damage to the submucosal layer.

### Accumulation of PAMPs for RNA and DNA-sensing pathways in HSV-2-infected vaginal epithelium

To formally demonstrate that the RLR-MAVS and cGAS-STNG pathways can be activated in the FGT, we looked for molecular signatures of activation of these two pathways. In HSV-2-infected cells, we observed capsid-free viral DNA using EdC-labelled virus and click chemistry (Fig. 3A) in agreement with previous observation for HSV-1 (*43*). Corroborating with this, we detected cGAMP in the mouse vagina after HSV-2 infection in WT but not *cGas^-/-^* mice (Fig. 3B). dsRNA can be detected with the antibody J2 (*16*). After genital HSV-2 infection, we observed emergence of a strong J2 signal in the vaginal epithelium (Fig. 3C), this was also seen in the vaginal organoids (Fig. 3D), and in human ectocervical explants (Fig. 3E, Fig S2A-B). The latter were permissive to HSV-2 infection, and responsive to agonists for the RLR-MAVS and cGAS-STNG pathways (Fig S2C-I), thus indicating that the findings in the mouse model recapitulate phenomena also observed in humans. As a possible source for dsRNA during HSV-2 infection we found that mitochondrial dsRNA could be immunoprecipitated from cytosolic extracts of HSV-2-infected VK2 human vaginal ECs (Fig. 3F-G, Fig S2J), which correlated with infection-induced mitochondrial dysfunction and alteration of the fumarate-Fh1 axis (Fig. 3H-J), previously reported to lead to activation RLR-MAVS by mitochondrial dsRNA (*22, 44*). Thus, the microenvironment in HSV-2-infected vaginal ECs accumulate the PAMPs enabling activation of innate RNA- and DNA-sensing pathways.

**Figure 3.**
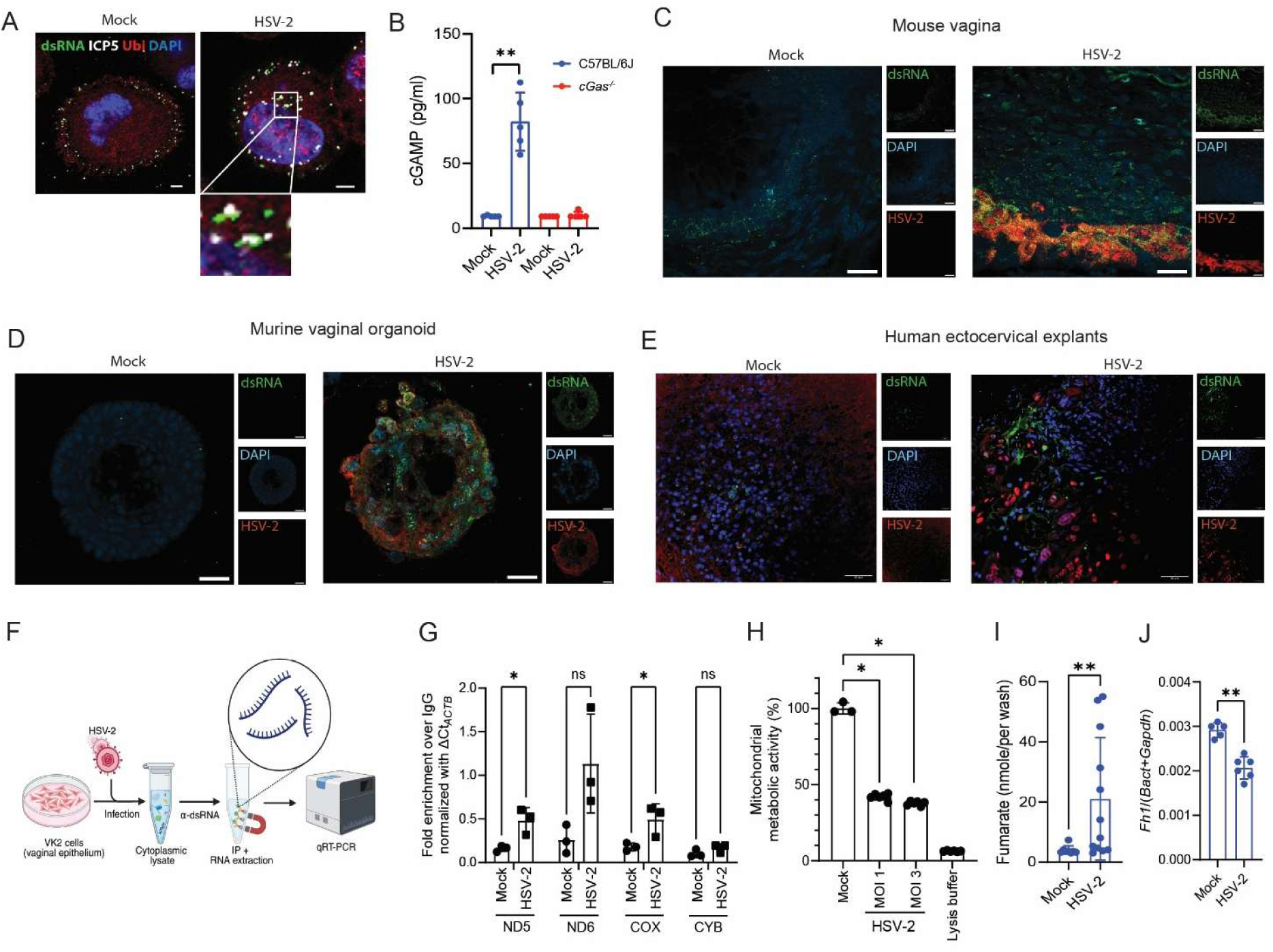
Accumulation of PAMPs for DNA and RNA-sensing pathways in HSV-2-infected epithelium. (A) HaCaT Cells were infected with EdC-labelled HSV-2 (MOI 1) for 1 h. Viral genomic DNA was visualized by click chemistry, and cells were stained with DAPI and antibodies against viral capsid major protein ICP5 and Ubiquitin. Scale bar: 10 μm. (B) Quantification of cGAMP in vaginal homogenates from WT and *cGas^-/-^* mice infected intravaginally with 6.7 × 10^4^ p.f.u. of HSV-2 for 48 h. (n = 5 mice/group). Representative confocal microscopy images of vaginal tissue sections from WT mice infected with 6.7 × 10^4^ p.f.u. of HSV-2 for 48 h (C), murine vaginal organoids infected with 2.5×10^5^ p.f.u HSV-2 for 48 h (D), and human ectocervical explants infected with 2×10^6^ p.f.u HSV-2 for 24 h (E), stained with anti-dsRNA (green), anti-HSV-2 (red), and DAPI (blue). (F) Illustration of the experimental procedure to examine for accumulation of dsRNA in the cytoplasm of HSV-2-infected cells. Created with Biorender. (G) Fold enrichment of mitochondrial RNAs in cytosolic anti-dsRNA antibody immunoprecipitates from cytotosolic lysates from VK2 cells infected for 24 h with HSV-2 (MOI 3) compared to mock. (H) Mitochondrial NADPH reductase activity of VK2 cells infected for 24 h with HSV-2 (MOI 1 and 3). (I) Fumarate levels in vaginal washes from C57BL/6J mice infected for 48 h with 6.7 × 10^4^ p.f.u. of HSV-2. (J) Levels of *Fh1* in RNA from vaginal organoids infected for 24 h with 2.5 × 10^5^ p.f.u. of HSV-2 normalized to average of *b-actin* and *Gapdh*. Data were shown as means +/- st.dev. (B, I, J) Wilcoxon rank-sum test. (G) ANOVA with a Turkey’s post hoc test. ^∗^*p* < 0.05, ^∗∗^*p* < 0.01; ns, not significant.

### Impaired TBK1 activation and control of HSV-2 in ECs lacking cGAS and RLR signaling

In order to start addressing the question as to how defective cGAS or RLR signaling impact on the local antiviral response in the vaginal microenvironment, we employed imaging mass cytometry (IMC) multiplex antibody staining together with GeoMx regional transcriptomics (Fig. 4A) on murine FGT samples isolated 24 and 48 h post infection. For the IMC, we used a panel of cell type markers combined with anti-pTBK1 to monitor PRR activation. We annotated 14 cell populations (Fig. 4B, Fig. S3A-B), including four epithelial subpopulations which exhibited distinct localization in the tissue (Fig. 4C). All epithelial cell populations showed elevated viral antigen load in *cGas^-/-^* and *Mavs^-/-^*mice compared to WT (Fig. 4D; Fig S3C). Looking across all annotated cell types, we observed the highest degree of TBK1 phosphorylation in ECs, arguing for a central role for ECs in evoking the early antiviral defense (Fig. 4E). pTBK1 was indeed induced in ECs upon HSV-2 infection in WT mice and was reduced in the ECs from *cGas^-/-^* and *Mavs^-/-^* (Fig 4F; Fig. S3D). The pTBK1+ signal was enriched in HSV-2^+^ versus HSV-2^-^ cells, suggesting PRR activation occurred to a higher degree in infected ECs (Fig. 4G). Using GeoMx regional transcriptomics, we next analyzed mucosal ECs and separated them into in HSV-2^+^ versus HSV-2^-^ cells (Fig. S3E). Interestingly, HSV-2 infection evoked an early ISG response in ECs from WT and *Mavs^-/-^* irrespective of whether they were HSV-2 positive or negative, but this was largely ablated in *cGas^-/-^* mice (Fig. 3H). Of note, later, on day 2 post infection the ISG response was partly induced in *cGas^-/-^* mice (Fig S3F), but the delayed response seemed not to protect against disease (Fig. 1B-C). The impaired expression of ISGs and chemokines in the FGT of *cGas^-/-^*mice was confirmed by RT-qPCR of vaginal tissue and ELISA of vaginal washes (Fig. 4I-M). In this material, we also confirmed the unaltered expression in *Mavs^-/-^*mice. In contrast, MAVS-deficiency did lead to modest reduction in ISG expression in vaginal organoids (Fig. 4N-Q), potentially explained by the more synchronized nature and high degree of productive infection in this material. Blocking STING signaling in *Mavs^-/-^* organoids nearly ablated HSV-2-induced ISG expression (Fig. S3G), suggesting these two pathways to be responsible for the bulk of the virus-induced response in ECs. Collectively, defective RNA and DNA sensing pathways lead to elevated HSV-2 replication in vaginal ECs, correlating with impaired activation of TBK1 in *cGas^-/-^* and *Mavs^-/-^*mice. The downstream antiviral IFN response is strongly impaired in *cGas^-/-^*ECs, but with only limited effect MAVS-deficient vagina ECs.

**Figure 4.**
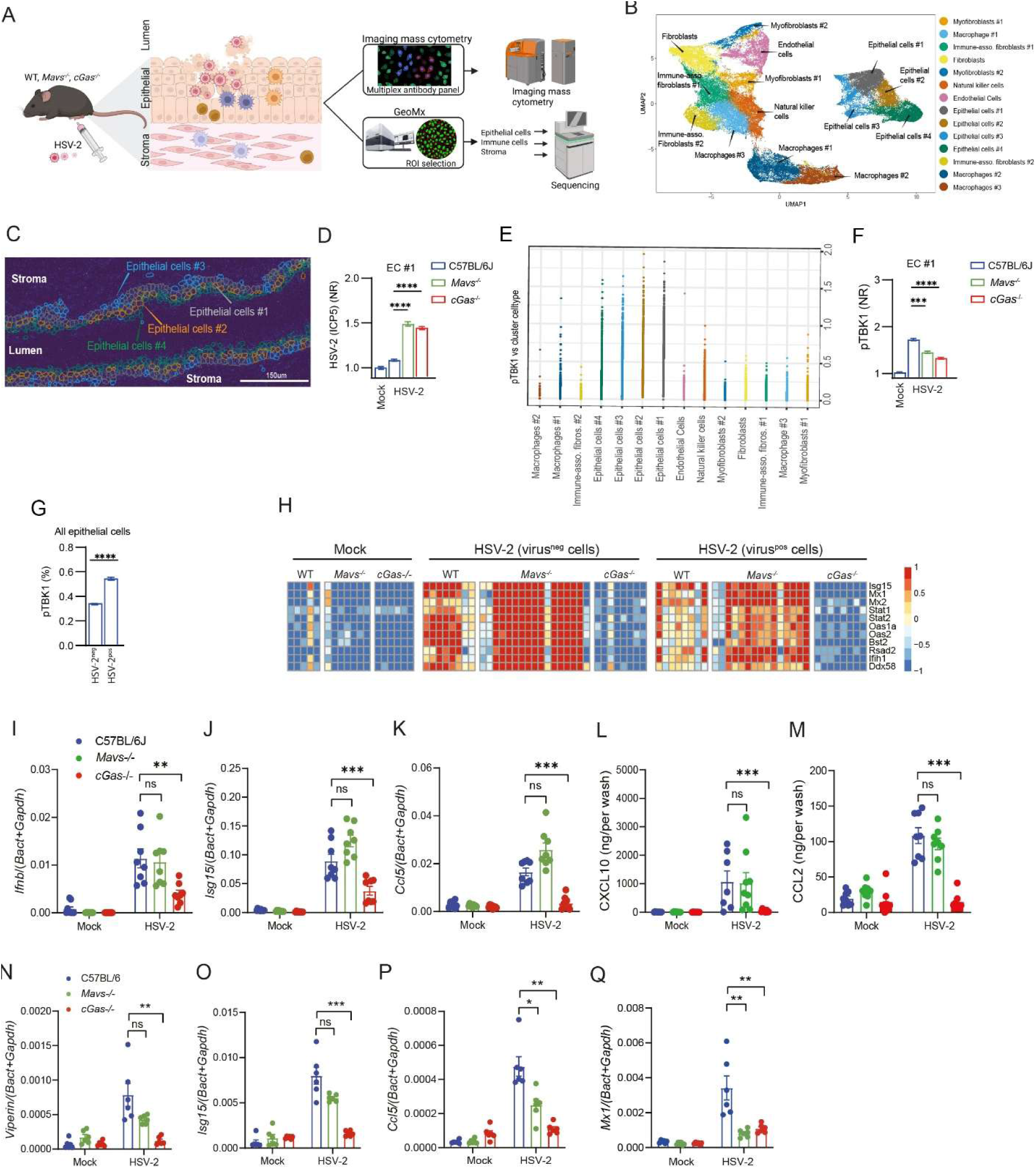
Early virus-induced expression of ISGs in vaginal epithelial cells is impaired in cGAS-deficient mice. (A) Illustration of experimental design and procedure for imaging mass cytometry and GeoMx analysis of HSV-2 infection in the vagina. The image was generated with Biorender. (B) UMAP of annotated cell populations in the imaging mass cytometry (IMC) dataset. (C) Representative IMC image with spatial overlay of epithelial subpopulations defined in (B). (D) Quantification of HSV-2 (ICP5) in epithelial cell (EC) population #1 in WT, *cGas^-/-^*, and *Mavs^-/-^* mice 48 h after infection (n= 780-2029 cells/group, from 3-9 ROIs). The values are normalized to WT-mock group (E) pTBK1 levels across annotated cell types. Each dot represents single cells. (F) Quantification of pTBK1 in epithelial cell (EC) population #1 in WT, *cGas^-/-^*, and *Mavs^-/-^* mice 24 h after infection with HSV-2 (n= 780-917 cells/group, from 3-9 ROIs). The values are normalized to WT-mock group (G) Percentage of pTBK1^+^ in HSV-2^+^ and HSV-2^-^ ECs (all EC subpopulations merged) 24 h after infection (n= 1529-1266 cells/group, from 3-9 ROIs). (H) Heatmap of selected ISG transcripts in ECs analyzed with GeoMx digital spatial transcript profiler. HSV-2^+^ and HSV-2^-^ ECs were isolated from the vagina of mice infected for 24 h. 83 CD11b^-^ AOIs in the epithelial layers were analyzed from 2-4 mice/group). (I-M) WT, *Mavs^-/-^*, and *cGas^-/-^* mice were infected for 48 h. Total RNA and vaginal washes were isolated and analyzed for the indicated transcripts (I-K) and cytokines (L-M), respectively. (n = 8 mice/group). (N-Q) Levels of *Viperin*, *Isg15*, *Ccl5,* and *Mx1* mRNA in vaginal organoids from WT*, cGas^-/-^*, and *Mavs^-/-^* mice 24 h after infection. Data were shown as means +/- st.dev. (D, F, I-Q) The p values were calculated using 2-way ANOVA with Bonferroni’s test. ^∗^*p* < 0.05, ^∗∗^*p* < 0.01; ^∗∗∗^*p* < 0.001; ^∗∗∗∗^*p* < 0.0001; ns, not significant. All mice in this figure were infected intravaginally with HSV-2 (6.7 × 10^4^ p.f.u. per mouse).

### Macrophages localize to foci of infection to interact with ECs

Along with the early production of ISGs by ECs, we observed influx of both myeloid and lymphoid cells into the FGT (Fig. S4A). While the influx of immune cells was apparent from day 2 in WT and *cGas^-/-^*mice, it was observed already on day 1 in *Mavs^-/-^*. We observed elevated viral load in the *Mavs^-/-^* mice already on day 1 (Fig. S4B), potentially triggering premature immune cell recruitment in the presence of an operative cGAS-STING pathway. We annotated three macrophage populations and one natural killer (NK) cell population (CD3^+^CD4^÷^CD8^÷^). While Macrophage #1 and #3 were observed in all mice, macrophage #2 accumulated in significant amounts on day 2 only in *cGas^-/-^* and *Mavs^-/-^* mice correlating with elevated viral load. This population was fibrinogen positive (Fig. S3B and Fig. S4A), suggesting an inflammation-associated phenotype.

When analyzing the IMC data we observed that macrophages #1 and #2 localized to the foci of virus infection, while macrophage #3 localized to the submucosal area (Fig. 5A). NK cells localized to both the infectious plaque and the submucosal area. Consistent with the plaque-associated location of macrophage #1, this cell population contained high levels of viral antigen across all genotypes correlating with viral load: WT < *Mavs^-/-^ < cGas^-/-^* (Fig. 5B). Examining for transcripts from the epithelium-localized CD11b+ cells (likely macrophage #1 and #2) by GeoMx, we found the ISG and chemokine levels to be highly elevated in infected *Mavs^-/-^* mice and moderately decreased in *cGas^-/-^* mice (Fig. 5C). In contrast, expression of the inflammatory cytokines *Tnf* and *Il1b* by these myeloid populations were elevated in both *cGas^-/-^* and *Mavs^-/-^* mice.

**Figure 5.**
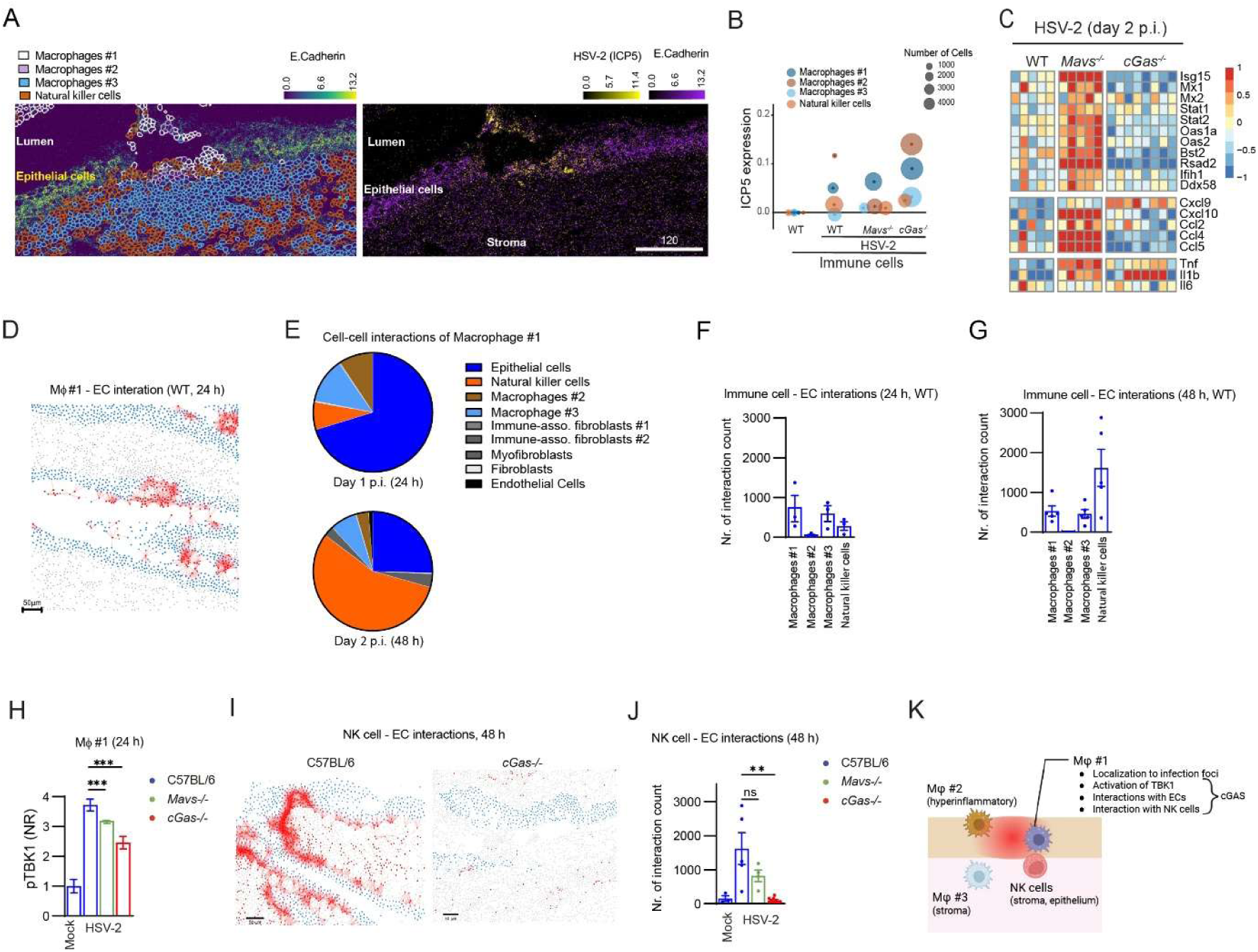
Impaired immune signaling in cGAS-deficient infection-foci-localized macrophages interacting with ECs. (A) Representative image of localizations of immune cell populations (left) and virus (right) in the vaginal mucosa of C57BL/6J mice infected for 48 h. (B) Bubble plot showing HSV-2 positive cells across immune cell populations in the vagina from WT*, cGas^-/-^*, and *Mavs^-/-^* mice infected for 48 h. (C) Heatmap of selected ISG, chemokine, and cytokine transcripts in CD11b^+^ cells analyzed with GeoMx digital spatial transcript profiler. Cells were isolated from the vagina of mice infected with HSV-2 for 48 h. 18 CD11b^+^ AOIs in the epithelial layers were analyze from 2-4 mice/group. (D) Representative image overlaid with spatial interaction analysis for macrophage #1 and ECs in the vagina of WT mice 24 h after vaginal infection. Spatial interaction analysis based on proximity (≤ 30 µm), Mj#1 (red), EC (blue), and other cells (gray) Scalebar, 50 mm. (E) Pie chart representation of non-homotypic cell-cell interactions of macrophage #1 in the vagina of WT mice on day 1 and 2 after infection with HSV-2. (F-G) Quantification of immune-cell - EC interactions in the vagina 24 h (F), and 48 h (G) after infection with HSV-2. (Each dot represents one ROI). (H) Quantification of pTBK1 in macrophage #1 in WT, *cGas^-/-^*, and *Mavs^-/-^*mice 24 h after infection with HSV-2. (n=5-1291 cells/group, from 3-9 ROIs). (I) Representative image overlaid with spatial interaction analysis for NK cells and ECs in the vagina of WT mice 48 h after infection. Spatial interaction analysis based on proximity (≤ 30 µm), NK cells (red), EC (blue), and other cells (gray) Scalebar, 50 mm. (J) Quantification of NK cell - EC interactions in the vagina of WT, *cGas^-/-^*, and *Mavs^-/-^* mice 48 h after HSV-2 infection (Each dot represents one ROI). (K) Summary of findings on localization and key functions of leukocyte populations in the HSV-2-infected vagina. (H) The p values were calculated using 2-way ANOVA with Bonferroni’s test. (J) *P* values were calculated using ANOVA with a Kruskal-Wallis multiple comparison test. ^∗^*p* < 0.05, ^∗∗^*p* < 0.01; ^∗∗∗^*p* < 0.001; ^∗∗∗∗^*p* < 0.0001; ns, not significant. All mice in this figure were infected intravaginally with HSV-2 (6.7 × 10^4^ p.f.u. per mouse).

We were particularly interested in the potential contribution of macrophage #1 to antiviral activity given its localization, and also the observation that it was the leukocyte population with highest degree of TBK1 activation (Fig. 4E). Cell-cell interaction analyses of the IMC dataset showed that this cell population interacted extensively with ECs from its mucosal localization (Fig. 5D-E, Fig. S4C), and was the first immune population to engage in interactions with ECs during infection (Fig 5F), while several cell populations were engaged on day 2 p.i. (Fig. 5G). Interestingly, macrophage #1 showed reduced levels of pTBK1 particularly in *cGas^-/-^* mice (Fig. 5H), but compatible interactions with ECs although delayed in *cGas^-/-^* mice (Fig. S4D-E). On day 2 p.i. macrophage #1 additionally interacted extensively with NK cells, which were recruited to interact with ECs (Fig. 5I). Of note, this interaction was largely ablated in *cGas^-/-^* mice (Fig. 5I-J), pointing towards a role for macrophage #1 in coordination of immune cell activity. Analysis of the interactions of macrophage #3 with ECs, showed that this cell population did interact with ECs at the stroma-epithelium interphase (Fig. S4F-G), but no difference between genotypes were observed for #3 (Fig. S4H). Finally, macrophage #2 interacted extensively with ECs on day 2 p.i. in both *cGas^-/-^* and *Mavs^-/-^* mice, having a pronounced luminal position (Fig. S4I-J). Collectively, immune cells are abundantly recruited to the FGT after genital HSV-2 at day 2 p.i. and we identify a macrophage population #1 that starts to accumulate on day 1 and localizes to foci of infection to activate TBK1 in a manner highly dependent on cGAS and also to engage in interactions with NK cells (Fig 5K). Elevated and sustained viral load in the vagina of *cGas^-/-^* and *Mavs^-/-^* mice may lead to development of the hyperinflammatory macrophage population #2.

### The antiviral action of cGAS and MAVS are exerted through different mechanisms

The GeoMx regional transcriptomics data strongly suggested that early expression of ISGs by ECs was centrally involved in the antiviral action of the cGAS-STNG pathway in genital herpes. To test this, we treated WT and *cGas^-/-^* mice intravaginally with IFNβ 12 h post HSV-2 infection and evaluated viral load in vaginal washes. While the viral load was much higher in the *cGas^-/-^*mice than the WT mice, the IFN treatment rescued the control of virus to the same level as observed in WT mice (Fig. 6A-B).

**Figure 6.**
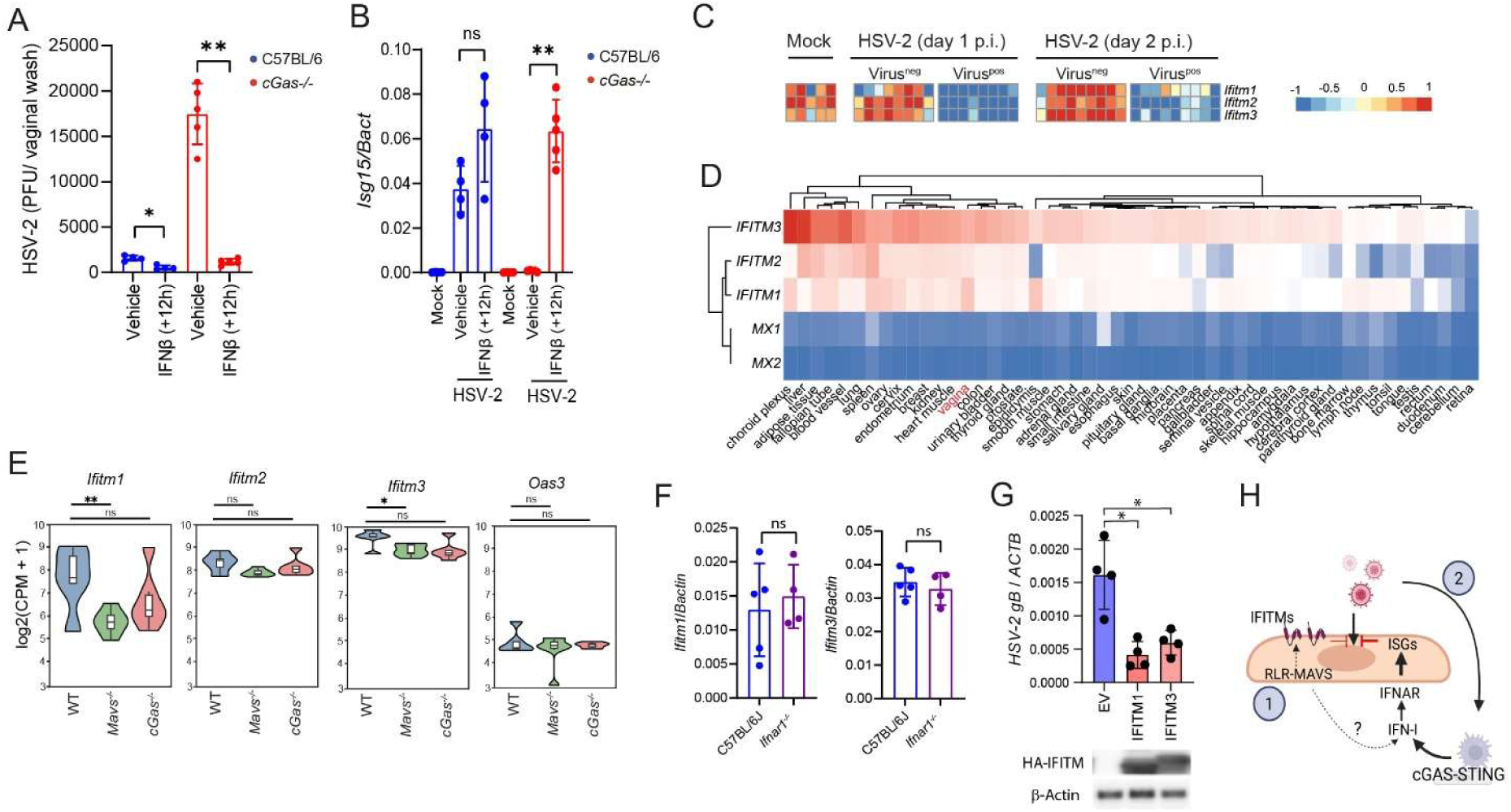
MAVS and cGAS utilize distinct mechanisms to restrict HSV-2 in the FGT. (A) WT and *cGas^-/-^* mice were infected intravaginally with HSV-2 (6.7 × 10^4^ p.f.u. per mouse) and treated locally with IFNb (10 mg) 12 hours later. Vaginal washes were isolated 48 h post infection and analyzed for viral load. (B) Mice were treated as in panel A and total vaginal RNA was isolated 24 h post infection, analyzed for *Isg15* and normalized to *Bactin*. (C) Heatmap of *Ifitm1*-*3* transcripts in WT ECs from mock- and HSV-2-infected mice (6.7 × 10^4^ p.f.u. per mouse, 24 h and 48 h) analyzed with GeoMx digital spatial transcript profiler. HSV-2^+^ and HSV-2^-^ ECs were isolated from the vagina of the mice (39 CD11b^-^ AOIs in the epithelial layers were analyze from 2-4 mice/group). (D) Heatmap of basal expression of *IFITM1-3*, and *MX1, 2* across human tissues. Data were obtained from the Human Protein Atlas Consensus dataset. (E) Violin blots of *Ifitm1, 2, 3,* and *Oas3* expression in vaginal ECs from uninfected WT*, cGas^-/-^*, and *Mavs^-/-^* mice. (F) *Ifitm1* and 3 expression in the vagina from uninfected WT and *Ifnar1^-/-^*, and *Mavs^-/-^* mice. (G) HSV-2 gB mRNA expression levels in human vaginal epithelial VK2 cells transfected with Empty vector (EV) or HA-IFITM1, 3. The cells were infected with HSV-2 (MOI 1) for 24 h. (n = 4 biological replicates). Transfections of ITITM1,3 are shown in the lower panel. (H) Summary of findings in the FGT. cGAS restricts HSV-2 through induction of IFN-ISGs, whereas MAVS, despite contributing only marginally to inducible IFN–ISG responses, is required for constitutive IFITM1 and IFITM3 expression in ECs and thus for the intrinsic barrier to HSV-2 infection. (A, B, G) Data were shown as means +/- st.dev. and analyzed using Wilcoxon rank-sum test. (E) DEGs were identified using generalized linear models based on a negative binomial distribution (DESeq2). Statistical significance was assessed via Wald test with Benjamini-Hochberg correction. Plots show means ± st.dev ^∗^*p* < 0.05, ^∗∗^*p* < 0.01; ns, not significant.

In the analysis of gene expression in vaginal ECs we noted that while the majority of ISGs were expressed at low levels at steady state, a small subset of genes normally annotated as ISGs were expressed at high basal levels in WT mice, notably the IFITMs (Fig. 6C). Mining human expression data showed that IFITMs are highly expressed constitutively across many human tissues, including the vagina (Fig. 6D). Interestingly, the basal expression of *Ifitm1*, and to a lesser extent *Ifitm3*, in mouse vaginal ECs was dependent on MAVS but not cGAS (Fig. 6E). The basal expression of *Ifitm1* and *3* in the mouse vagina was independent on IFNAR1 suggesting MAVS to drive *Ifitm* expression in vaginal ECs independently of intermediary IFN-I gene expression (Fig. 6F). To examine whether the high basal expression of IFITM1 and 3 impacts on restriction of HSV-2, we overexpressed the genes in human vaginal epithelial VK2 cells and infected with HSV-2. Interestingly, elevated basal expression of IFITM1 and 3 did indeed reduce HSV-2 replication in vaginal ECs (Fig. 6G). Collectively, these data suggest that while cGAS exerts its anti-HSV-2 activity through induced IFN-ISG expression, MAVS may have a minor role in induced IFN-ISG expression but is essential for constitutive expression of IFITM1, 3 in ECs to impose a barrier for establishment of HSV-2 infection (Fig, 6H).

### cGAS deficiency enables viral access to peripheral neurons and spread to the CNS

Given the impairment of the antiviral activity by ECs in *cGas^-/-^*and *Mavs-/-* mice, we were interested in knowing whether this impacted on viral spread to neuronal tissue. Analysis of the IMC data showed that while the majority of cell types remained HSV-2 negative during infection in WT and *Mavs^-/-^* mice, except ECs and some infiltrating immune cells, largely all cell types became positive on day 2 post infection in *cGas^-/-^*mice (Fig. 7A). This included fibroblasts, which localize to the submucosal area. The deeper penetration of virus into the tissue in *cGas^-/-^* mice was in agreement with the stainings of vaginal tissue sections for HSV-2 (Fig. 2). Interestingly, the ISG response was observed to extend into the stroma in WT and *Mavs^-/-^*, but not *cGas^-/-^* mice (Fig. 7B). These data raised the question as to whether cGAS-deficiency facilitated access to tissue-intervening neurons. Indeed, co-staining of stromal tissue from infected mice with HSV-2 and the neuronal marker Beta III tubulin showed high abundance of virus around neurons in the stroma of *cGas^-/-^* but not *Mavs-/-* or WT mice (Fig. 7C). DRGs contain the cell bodies of sensory neurons that innervate the vagina and connect to the spinal cord. When isolating DRG and spinal cord from infected mice, we observed significantly higher viral gene expression in PNS and CNS neurons from *cGas^-/-^* compared to WT mice (Fig. 7D-F). Collectively, while both the cGAS-STING and RLR-MAVS pathway exert control of HSV-2 infection in the vaginal epithelium, the cGAS-STING pathway additionally limits virus spread and hence neuroinvasion.

**Figure 7.**
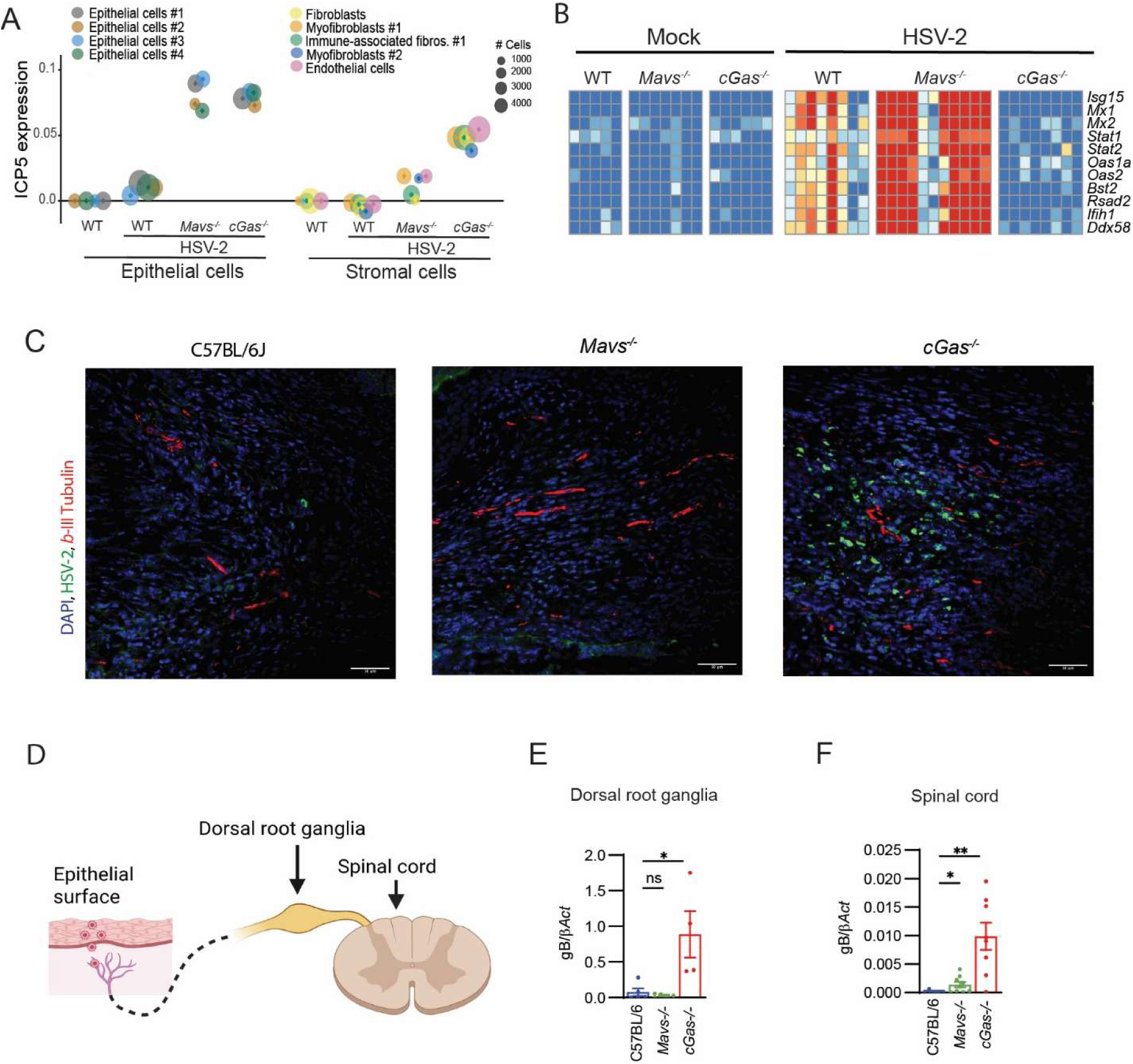
cGAS deficiency enables viral access to peripheral neurons and spread to the CNS. (A) Bubble plot showing HSV-2 positive cells across epithelial and stromal cell populations in the vagina from WT*, cGas^-/-^*, and *Mavs^-/-^* mice infected for 48 h.. (B) Heatmap of selected ISG transcripts in stroma analyzed with GeoMx digital spatial transcript profiler. The stroma was isolated from the vagina of mice infected for 24 h. (45 ROIs in the stroma layers were analyze from 2-4 mice/group)). (C) Immunofluorescence staining of stroma areas of mouse vagina after 48 h with HSV-2 (green), and neuronal marker, Beta III-tubulin (red). Scale bar: 50 μm. (D) Illustration of how viral spread to the submucosal area enabling may facilitate infections of neurons and spread to the dorsal root ganglia and spinal cord. Image made in Biorender. (E-F) Dorsal root ganglia and spinal cord were isolated from WT*, cGas^-/-^*, and *Mavs^-/-^* mice infected for 5 days with HSV-2, and viral *gB* transcripts were quantified by RT-qPCR and shown as normalization to *Bactin*. Data were shown as means +/- st.dev. ANOVA with a Turkey’s post hoc test. ^∗^*p* < 0.05, ^∗∗^*p* < 0.01. All mice in this figure were infected intravaginally with HSV-2 (6.7 × 10^4^ p.f.u. per mouse).

## DISCUSSION

Genital herpes is a prevalent sexually transmitted infection globally (*1*). It is characterized by painful sores in the genital area, with HSV-2 being the most common cause of genital herpes. Moreover, HSV-2 can spread from ECs to intervening peripheral neurons to establish latent infections that can periodically reactivate. This can be particularly critical during vaginal childbirth with high risk of transmission and life-threatening disseminated disease in the newborn. HSV-2 can spread to the CNS in adults giving rise to recurrent meningitis. Here we report that innate immunological nucleic acid-sensing pathways play distinct roles in exerting antiviral activity in the FGT. While the cGAS-STING and RLR-MAVS pathways cooperate to control HSV-2 infection in the vaginal epithelium, only the cGAS-STING pathway is critically important for prevention of viral spread from ECs to intervening DRG neurons and the spinal cord. The two pathways employ different antiviral mechanisms. RLR-MAVS maintains basal expression of the antiviral ISG IFITM1 and also enables sensing of mitochondrial stress in productively infected ECs. In contrast, cGAS drives the induced type I IFN response across cell types and coordinates the early recruitment and activation of immune cells to the mucosa. The cGAS-dependent IFN response extends beyond the infected ECs and into the stroma to protect from viral spread. Our work demonstrates how distinct pathogen-sensing pathways cooperate to protect the infected tissue.

A central observation of the present work is that *cGas^-/-^*mice exhibited a very strong vaginal HSV-2 infection phenotype, *Mavs^-/-^*showed a clear but intermediate phenotype, while *Tlr3^-/-^*mice did not have a phenotype at low infection doses in this model. These data suggest non-redundant roles for DNA and RNA sensing pathways in control of HSV-2 in the FGT. Ocular HSV-2 infection showed the same effects, potentially suggesting that the dual involvement of the cGAS-STING and the RLR-MAVS pathways in control of HSV-2 infections may be a general phenomenon across different portals of entry for HSV infections. Further studies are needed to explore whether this is also applies to infections with other DNA viruses, several of which have been reported to be able to activate both RNA and DNA sensing pathways (*17, 18, 23–25*). For RNA viruses, there is a large array of examples of non-redundant roles for RNA and DNA sensing pathways in antiviral defense (*27, 45–48*), with the viral genome being the prime RNA PAMP, and mitochondrial DNA the major DNA PAMP (*26, 49–53*). For example, in the case of paramyxoviruses, RNA sensors have been reported to be insufficient for the effective control of infection *in vivo*, and the cGAS-STING pathway is required for full IFN response and antiviral defense (*47*). In this work we find that productive HSV-2 infection in vaginal ECs leads to release of mitochondrial dsRNA into the cytoplasm, and also that a subset of ISGs are constitutively expressed in a MAVS dependent manner, potentially suggesting low-grade RNA-sensing during steady state. For the cGAS-STING pathway, we detected HSV-2 genomic DNA in the cytoplasm of infected cells suggesting that viral genomic DNA directly triggers the cGAS-STING pathway. The present work therefore provides mechanistic information on how the involvement of multiple nucleic acid pathways enables the organism to impose a basal barrier against infection as well as to sense both viral genomes and virus-induced activities. This allows activation of a broader and more robust immune response and likely also more difficult to evade by viruses.

At the level of pathology, we observed that the focal epithelial cell death observed in the infected WT mice was significantly expanded in *Mavs^-/-^*mice, but still retained within the epithelial cell layer. By contrast, in *cGas^-/-^*mice, the extensive cell death in ECs was accompanied by deeper progression of the necrotic response into the submucosal stromal layer. This suggested that *cGas^-/-^* and *Mavs^-/-^* mice exert qualitatively - and not only quantitatively – different anti-HSV-2 responses. In line with this, cGAS-deficiency largely ablated the early ISG response in vaginal ECs irrespective of whether the cells were infected or not. Intravaginal treatment of cGAS-deficient mice with IFNβ 12 h after infection rescued the antiviral response. Since pTBK1 was mainly observed in HSV-2 positive cells, this suggests that productively infected cells rapidly induce type I IFN gene expression in a cGAS-dependent manner, which leads to ISG expression across the epithelium to impose an antiviral state. With respect to MAVS, we observed a modest effect on ISG expression in organoids, which was not seen in the more complex in vivo situation, including for instance infiltration of immune cells. However, we found that IFITM1 and 3 were expressed at high basal levels in vaginal ECs in a MAVS-dependent manner, were not induced upon infection, and were in fact downregulated in HSV-2-infected cells. IFITMs are IFN-induced in many cell types, where they localize to membrane to block viral entry, thereby restricting diverse enveloped viruses across multiple cell types (*54*). Overexpression of IFITM1 led to reduced replication of HSV-2. IFITM1 and 3 have previously been suggested to restrict HSV-1 infection (*55, 56*), and we suggest that MAVS-IFN-dependent constitutive expression of these two ISGs in vaginal ECs imposes a barrier for HSV-2 and potentially other viruses. Future studies should explore the mechanism through which HSV-2 down-regulates IFITM expression, which may be important for establishing infection at mucosal surfaces.

Another important observation from this study is that the impaired IFN response in *cGas^-/-^*mice is found only at the earliest time points, while it is largely indistinguishable from infected WT mice later. The delayed response is likely mediated by RLR-MAVS signaling, since productive HSV-2 infection leads to release of dsRNA into the cytoplasm and the blockage of STING in MAVS-deficient organoids ablated ISG expression. Despite the eventual rescue of the IFN response in *cGas^-/-^* mice, the elevated susceptibility to HSV-2 infection is not transient and does lead to high lethality. These data show the importance of timing when it comes to IFN responses. If the type I IFN response is not evoked very early after acute infection, its antiviral activity is much less potent, and it may even amplify inflammation. In fact, we do indeed find augmented inflammatory gene expression in the *cGas^-/-^* mice on day 2 and also elevated levels of fibrinogen positive macrophages. The phenomenon of delayed IFN responses being a major factor conferring susceptibility to viral infections and amplification of immunopathology has also been observed in other viral infections, including SARS-CoV infections (*57*).

We observed infiltration of immune cells into the infected FGT. This included three macrophage populations, and one leukocyte population annotated NK cells (CD3^+^CD4^-^CD8^-^). At this stage we cannot distinguish whether the three macrophage populations are distinct or rather represent different stages of the same infiltrating population: #3 (submucocal) -> #1 (infection foci, early) -> #2 (infection foci, late). Although more analysis is required to clarify this, the authors favor the latter explanation. Macrophage population #1 (CD45^+^CD11b^+^F4/80^+^) localized to the foci of infection, exhibited the highest level of pTBK1 across the leukocytes, interacted extensively with ECs, and later also and NK cells, which are important for control of genital herpes (*13*). The interaction of macrophage #1 with ECs was delayed in *cGas^-/-^* mice, but comparable across genotypes on day 2. However, this population showed lower levels of pTBK1, particularly in *cGas^-/-^* mice. These data suggest that macrophage #1 is recruited to the foci of HSV-2 infection to orchestrate antiviral activities in a cGAS-dependent manner. We propose that extensive or prolonged exposure of macrophage #1 to virus trigger a hyperactivation stage, #1 -> #2, with expression of fibronectin and the inflammatory cytokines *Tnf* and *IL1b* as observed on day 2 in *cGas^-/-^* and *Mavs^-/-^*mice. Therefore, altering the tissue-intrinsic early antiviral activity of the epithelium changes the nature of the activity of infiltrating leukocytes. This is driven by both higher viral load, alterations in the cytokine microenvironment, and extensive cell death.

Finally, we observed that cGAS restricts HSV-2 access to the neurons intervening the vaginal tissue, and downstream viral access to the CNS. While papers from several laboratories, including our own, have shown a role for RNA and DNA sensing innate immune pathway in antiviral activity in the CNS (*30, 45, 58*), this is to the best of our knowledge the first report of a PRR restricting a neurotropic virus to gain access to peripheral neurons from the primary site of infection. Mechanistically, we show that loss of cGAS leads to impairment of the ISG response to extend into the submucosal stroma. We propose this facilitates HSV-2 spread to the submucosal area, where the virus can infect intervening sensory neurons. We previously reported a patient with seven confirmed cases of HSV-2 meningitis, who carried a heterozygous variant of *IKBKE* exerting dominant negative activity of both IKKε and TBK1 (*59*). Cells from the patient exhibited functional defect of the cGAS-STING pathway. Together with the results from the present work, these studies propose an important role for the cGAS-STING pathway in preventing CNS diseases by HSV-2, as it occurs in recurring HSV-2 meningitis.

In summary, this study identifies distinct and non-redundant roles of the RLRs-MAVS and cGAS-STING pathways in control of genital HSV-2 infection. The findings in the mouse model were recapitulated also in a human genital explants model. Moreover, we report a distinct role for the cGAS-STING pathway in limiting viral spread beyond the epithelium and into intervening neurons. Although all pathogens activate multiple PRRs, the mechanisms through which PRRs cooperate to limit infections are poorly understood despite the obvious importance. This study highlights the complex nature of the cooperative orchestration of antiviral responses, which are finely tuned to balance antiviral defense against immunopathology. Understanding how immune mechanisms cooperate and cross-talk will advance knowledge on disease pathogenesis and device improved treatment strategies.

## METHODS

### Mice

Mice used in this study included age-matched (6–10 weeks of age) C57BL/6J wild-type, *cGas^-/-^* (C57BL/6-Mb21d1tm1d(EUCOMM)Hmgu), *Tlr3^-/-^* (C57BL/6-Tlr3tm1Fl), and *Mavs^-/-^*(C57BL/6-Mavstm1Tsc). C57BL/6J, *cGas^-/-^ Mavs^-/-^*, and *Tlr3^-/-^* mice were bred at Taconic M&B, and Aarhus University. All experiments were conducted at Aarhus University and have been reviewed and approved by Danish government authorities, complying with Danish laws. Measures were taken to minimize animal suffering, with daily monitoring post-infection. The mice were not randomized but after HSV infection, the information about mice strain and treatment were blinded to the investigators. No animals were excluded from the analysis.

### Murine HSV infection model

The murine HSV ocular infection model mimics the potential and natural route of HSV entry in HSE. Male mice were anesthetized with intraperitoneal injection of Ketamine (100 mg kg^-1^ body weight) and xylazine (10 mg kg^-1^ body weight). Corneas were sacrificed in a 10 × 10 crosshatch pattern and mice were inoculated with either 1 × 10^6^ PFU HSV-1 or 200 PFU HSV-2 in 5ul of infection medium (DMEM containing 200 IU ml ^-1^ penicillin and 200mg ml ^-1^ streptomycin). Mice were killed once they met endpoint criteria or at the specified times post infection. The murine HSV vaginal infection model mimics the natural route of HSV entry in genital herpes. Female mice were pretreated with subcutaneous injection of 2mg Depo-Provera, to induce the diestrus stage and increase susceptibility to vaginal HSV-2 infection (*60*). Six days later, the mice were anesthetized and inoculated intravaginally with 6.7 × 10^4^ PFU HSV-2 or 6.7 × 10^6^ PFU HSV-1 (Mckrae strain). The mice were then placed on their backs for 10 min. In the survival experiments, infected mice were monitored for disease symptoms daily and euthanized when they reached human endpoints: weight loss >20%, severe inflammation with ulceration in the genitoanal region, paresis of the hind limbs, or hunchback with antisocial behavior and facial expressions of pain.

### Isolation of DRGs and spinal cord

Spinal cords were harvested from euthanized mice immediately postmortem. Following exposure of the vertebral column, the spinal column was excised and placed in ice-cold buffered solution. The spinal cord was isolated by opening the vertebral canal and gently releasing the cord from the surrounding meninges and nerve roots. The isolated spinal cord was briefly rinsed in cold buffer to remove blood and debris and then processed for downstream analysis. To isolate L5, L6 and S1 DRGs, mice were transcardially perfused with PBS and the spinal canal was exposed by cutting the spinal vertebrae bilaterally at each spinal segment. Individual DRGs was identified by counting from costae and carefully excised with minimal attached nerve tissue and collected into ice-cold buffer (*61*). Where indicated, DRGs were pooled by animal and/or by spinal level (e.g., cervical, thoracic, lumbar, sacral) prior to downstream processing. All instruments and collection surfaces were kept cold to preserve tissue integrity.

### Human ectocervical explants

Uterine specimens were obtained from patients undergoing hysterectomy for benign conditions, such as endometriosis, adenomyosis, or benign myoma, at the Danderyd Hospital, Sweden. Only cases with normal cervical histology based on routine pathological examination were included. Exclusion criteria were: clinical symptoms of sexually transmitted infections, systemic immunosuppressive therapy, and past or present human papillomavirus (HPV)-related cervical lesions. A sample of mucosa was dissected into approximately 3 × 3 × 3 mm ectocervical tissue blocks (“explants”) within 2 h from surgery. The explants were placed in a 12-well plate (one explant per well) in culture medium prepared by supplementing RPMI 1640 with gentamicin 50μg/ml, Fungizone 2.5μg/ml, non-essential amino acids, and sodium pyruvate 1mM (all from ThermoFisher Scientific). Each well was supplied with either 100ug cGAMP (InvivoGen), 70ng RIG-I agonist (M8) transfected with Lipofectamine™ RNAiMAX Transfection Reagent (ThermoFisher Scientific), or 2*10E6 PFU HSV-2, followed by incubation at 37°C, CO_2_ 5%, and humidity 95% for 24 hrs.

### Murine vaginal organoid

Organoids were generated as described (*62*). The vaginas of 6-10 weeks old female mice were dissected and washed with Ca^2+^ and Mg^2+^ free Hanks Balanced Salt Solution. The tissues were placed in DMEM/F12 medium supplemented with 10% fetal bovine serum. The epithelium was then peeled off with the help of a pair of forceps. Following two successive washes in DMEM/F12, the single-cell suspension was prepared using 40mm cell strainer before plating. For organoid culture, 20,000 cells were resuspended in 50ul Matrigel and cultured in DMEM/F12 supplemented with 2% Ultraserum-G, 1% Penicillin-Streptomycin, 50ng/ml mouse EGF, 0.5mM A83-01 (TGFb/Alk inhibitor) and 10mM Y-27632 dihydrochloride (ROCK inhibitor). The medium was changed every 48 h.

### Reverse transcription quantitative PCR

Vaginas, brain stems, eyeballs and spinal cords from mice or human ectocervical explants were homogenized in PBS. RNA from brain stems, eyeballs, and spinal cords was extracted through High Pure RNA Isolation Kit (Roche), while RNA from vaginas and human ectocervical explants was isolated through Mini kit for RNA purification (MACHEREY-NAGEL). Gene expression was determined by reverse transcriptase quantitative PCR (RT-qPCR), using TaqMan (Applied Biosystems) systems. 100ng of RNA was used for each reaction. Expression levels were quantified relative to the expression of β-Actin.

### Immunohistochemistry

Mice were perfused prior to tissue collection. Dissected vaginal tissues from mice or human ectocervical explants were fixed in 4% formaldehyde, embedded in paraffin, and sectioned at a thickness of 4 µm. Sections were subjected to antigen retrieval in citrate buffer, followed by blocking with 5% bovine serum albumin (BSA) to reduce nonspecific binding.

Sections were incubated overnight at 4 °C in a humidified chamber with the following primary antibodies diluted in blocking buffer: anti–cytokeratin 5 (K5; 1:400; ab52635; Abcam), anti–HSV-1/2 (1:500; B0116; Dako or ab6508; Abcam), anti–double-stranded RNA (dsRNA) clone J2 (1:400; MABE11134; Sigma-Aldrich), and anti–β3-tubulin (1:400; Cell Signaling Technology). Following day, sections were washed in TBS + 0.3% BSA and incubated for 1 h at room temperature with species-appropriate Alexa Fluor 488, Alexa Fluor 568 or Alexa Fluor 647–conjugated secondary antibodies (Thermo Fischer Scientific) diluted 1:400 in TBS + 1% BSA. Nuclei were counterstained with DAPI. Slides were mounted with antifade mounting medium and coverslipped prior to imaging.

Fluorescent images were acquired using a Zeiss LSM800 confocal microscope equipped with a 40× objective under identical acquisition settings for all experimental groups. Laser power, detector gain, and pinhole size were kept constant across samples to enable quantitative comparison. For each group, tissues from n = 4 animals were analyzed, with two nonadjacent sections examined per animal. Images were processed using Fiji (v2.1.0/1.53c) for figure preparation only. Brightness and contrast adjustments were applied uniformly across entire images and identically across experimental groups.

### Quantification of Fumarate

Vaginal washes were collected after 48 h p.i. by washing with 2 × 40 μL of DMEM and dilution to a final volume of 250 μL. Fumarate levels in the vaginal washes were measured using the Fumarate Detection Kit (ab102516; Abcam).

### Flow cytometry

Twenty microliters of propidium iodide (PI; 10 μg/mL; MedChemExpress, HY-D0815) were administered intravaginally for 10 min. Vaginal washes were collected as described above and stained with 1 μg/mL Hoechst 33342 (62249, Thermo Scientific) for 10 min at RT in the dark. Samples were filtered through a 70 μm polystyrene cell-strainer cap (352235, Falcon) washed with PBS to remove debris, and equal fraction of the virginal washes were subsequently used on a NovoCyte flow cytometer (ACEA Biosciences) and analyzed on NovoExpress software (both Agilent, Santa Clara, CA). The nr of dead (PI^+^) cells among total Hoechst^+^ cells was calculated and plotted.

### Click Chemistry

Propagation of EdC-labeled HSV-2 in BHK-21 cells and visualization of HSV-1 genome DNA by click chemistry was performed as described previously (*63*). In brief, cells were infected with EdC-HSV-2 at an MOI of 10 and fixed at the indicated time points postinfection by methanol at −20°C for 5 min. EdC-incorporated viral DNA was labeled using the Click-iT Alexa Fluor 488 Imaging Kit (C10337; Thermo Fisher Scientific) according to the manufacturer’s protocol. After washing away click reagent, cells were blocked with 1% BSA and subsequently stained with the indicated primary and secondary antibodies. Images were acquired on a Zeiss LSM 810 confocal microscope, using a 63 × 1.4 oil-immersion objective, and imaging processing was performed using Zen software (Zeiss).

### Imaging mass cytometry

Antibody-metal labeling for IMC: Antibodies compatible with immunocytochemistry that were BSA-free and at 100 µg quantity with at least 0.5 mg/ml concentration were purchased from different vendors (Table S1). Metal labeling of antibodies was performed using Maxpar® X8 Multimetal Labeling Kit as described by the manufacturer’s protocol (Standard BioTools).

FFPE slide tissue staining for IMC: FFPE fixed tissue slides were stained according to the standard protocol for FFPE sections (StandardBioTools PN 400322 04 PROTOCOL with minor modifications. FFPE Slides were baked for 30min at 80°C, dewaxed in Xyelene, Hydrated using sequentially, 99%,96%, 80%, 70% EtOH, and washed with PBS. Antigen retrival reagent pH9 (Agilent) incubation was done for 30min at 80°C. Slides were blocked with 3% BSA in PBS for 1 h and stained with a mixture of metal-labeled antibodies (Table S1) overnight at 40C in a hydration chamber. After staining, the slides were washed with TritonX-100 (0,2% in PBS) and PBS, and counterstained with Cell-ID™ Ir191/193 genomic DNA intercalator (Standard BioTools) at 1:400 dilution for 30min. Finally, the slides were washed with PBS followed by quick dip in ddH2O to remove the salts, dried and stored until ablated with the Hyperion imaging system.

IMC sample acquisition: Regions of Interest (ROI) were acquired by laser ablation with the Hyperion Imaging System (Standard BioTools) connected to a Helios CyTOF system (Standard BioTools). The instrument was tuned with 3-element tuning slides provided by the manufacturer prior to sample acquisitions. A minimum dual count of Lu 800, transient crosstalk <25% and resolution (first and second mass) >400 criteria were used to successfully pass the tuning of an instrument, as recommended by the manufacturer (Standard BioTools). ROIs of varies sizes were ablated, guided by HSV stained IHC adjacent serial sections, investigated for the presence of virus. For uninfected slides, a similar vaginal region was chosen. The data were obtained in .mcd (Standard BioTools standard file format) and .txt format.

IMC Data Processing and Analysis: Software packages Steinbock and IMC RTools (*64*) were used for data processing and analysis. Using Steinbock, .txt format datafiles were processed to generate images (.tiff) datafiles, including HotPixelFiltering (hpf50). In cases of corrupted data files (due to acquisition errors), .txt files were repaired by manual editing of the corrupted lines before Steinbock processing. Some images were cropped into sub-images, using ImageJ/Fuji software after HotPixelFiltering. Steinbock Software was further used to generate outputs such as segmentation (mesmer –minmax), quantification (intensities, regionprops) and neighborhood (neighbors --type expansion --dmax 4) data.

Subsequent Data Analysis was done using RStudio and the IMC RTools script with modifications (*64*). Data analysis steps included: Batch effect correction, Clustering, manual classification/annotation based on marker expression, single-cell visualization, image visualization, and border-based spatial interaction analysis. See Table S2 for detailed packages list.

### GeoMx Digital Spatial Profiling

The tissue was prepared and cut into 5 µm sections as described under Immunohistochemisty above. The protocol for GeoMx was followed as described previously with few modifications (*65*). Most importantly, antigen retrieval was performed at 99°C for 20 min, the concentration of proteinase K (Invitrogen cat. nr. AM2546) used was 0.8 µg/mL, and DSP RNA whole transcriptome detection probes used were GeoMx Mouse Whole Transcriptome v2 (WTA) (Bruker Spatial Biology). The primary antibodies used were CD11b (Biolegend, clone M1/70, cat. nr. 101204) in dilution 1:125 and HSV1/2 (Dako, Rabbit-polyclonal, GA521). The secondary antibodies and nuclear stains were done by using Donkey anti-rabbit AF488 (Invitrogen, A21206) in dilution 1:400 and Donkey anti-rat AF647 (Invitrogen, A78947) in dilution 1:400, (Syto83 Thermo Fisher, S11364) 400 nM.

ROI Selection Strategy: The slides were loaded onto the GeoMx DSP instrument (Bruker Spatial Biology) and scanned to produce a digital image of the tissue sections. The digital image was used to visualize tissue morphology based on the fluorescent labelled antibodies CD11b, HSV1/2 and the DNA stain SYTO 83. Regions of interest (ROI) were selected based on high morphological expression of HSV-positive areas. Epithelial and stroma was distinguished based on morphology and SYTO 83 staining. Multiple ROIs of various sizes (up to maximum ROI size of 660 × 785 µm) and with varying cell counts were selected.

ROI Segmentation/AOI Profiling and AOI Collection: After ROIs within epithelial layer were selected, the image analysis software integrated on the GeoMx DSP instrument was used to carry out threshold-based segmentation, to divide each ROI into distinct segments. The stroma and mock infected ROIs were not segmented further since they did not contain enough HSV^+^ or CD11b^+^ cells. However, the epithelial layer was segmented into individual biological compartments based on tissue morphology, CD11b, HSV expression, and cell count.

Each ROI was divided into 3 segments containing individual cell populations with CD11b^+^ HSV^+^, CD11b^-^ HSV^+^, CD11b^-^ HSV^-^. Due to the low abundance of CD11b^+^ HSV^-^ cells within the epithelial ROI, these cells were not collected. Each segment within a ROI corresponds to one area of illumination (AOI). 2-4 mice/ group was used. 74 ROIs from stroma, 36 CD11b^+^ AOIs in the epithelial layer, 135 CD11b^-^ AOIs in the epithelial layers were analyzed. Libraries were prepared according to the protocol described in (65) and sequenced on an Illumina NovaSeq 6000 platform (Illumina).

GeoMx data analysis: The generated FASTQ files were converted to output DCC files using the GeoMx NGS Pipeline v 2.3.3.10. Briefly, raw reads were trimmed to remove adapter sequences and overlapping paired-end reads were merged to reconstruct probe sequences containing barcode and UMI regions. The stitched reads were matched to probe barcode sequences in the reference assay to assign reads to target transcripts. PCR duplicates were removed using UMI sequences, resulting in deduplicated digital counts. Probe level counts were converted to gene level counts with the R package standR (v1.6.0.2). Initial gene-level filtering removed consistently low-expressing genes using function with addPerROIQC function with default parameters. Specifically, genes were excluded if their log-transformed counts per million fell below a dynamic threshold derived from a minimal count of 5 and the median library sizes in 90% of the ROIs. Subsequently, ROI-level quality control was conducted by excluding segments with fewer than 10 nuclei or a library size below 30,000 counts. For exploratory visualization, retained counts were normalized using the TMM method. Finally, differential expression analysis was performed on the raw counts using DESeq2, with the statistical model accounting for the specified experimental grouping variable and relevant covariates.

### Tissue-level gene expression profiling

To evaluate tissue-specific expression profiles, RNA expression data were acquired from the Human Protein Atlas (HPA) Consensus dataset. This resource summarizes gene-level expression across human tissues based on transcriptomic data from the HPA and GTEx cohorts. The acquired dataset was subsequently utilized to construct a heatmap, illustrating the tissue-wise expression distribution of the selected genes.

### Cytokine and cGAMP ELISAs

Cytokine/chemokine concentrations were quantified by ELISA. Murine CXCL10 and CCL5 were measured using commercially available ELISA kits (Bio-Techne) according to the manufacturer’s instructions. cGAMP levels were determined using a competitive ELISA kit (Cayman Chemical) following the manufacturer’s protocol. Absorbance was recorded on a microplate reader, and analyte concentrations were interpolated from standard curves generated with kit standards. Samples were analyzed in technical duplicates.

### Transfecton of VK2 cells with IFITMs

5×10^4^ VK2 cells in a 24-well plate were transfected with empty vector or pCMV-HA-IFITM using Lipofectamine 2000 (Thermo Fisher Scientific, 11668019). After 24 hours, the cells were infected with HSV-2 at MOI 3 for 6 hours. For WB, the cells were lysed with RIPA buffer supplemented with Benzonase (Sigma-Aldrich, E1014-25KU), cOmplete protease inhibitor cocktail (Roche, 5892953001) and 10 mM NaF (Sigma-Aldrich, S7920-100G). For RT-qPCR, the total RNAs were isolated using High Pure RNA isolation kit. The RNA levels of the genes were analyzed by RT-qPCR using TaqMan RNA to CT One Step Kit (Thermo Fisher Scientific, 4392938). Expression levels were determined by immunoblotting. Antibodies: Anti-β-Actin (Cell Signaling Technology, 8457S), anti-HA-Tag (Cell Signaling Technology, 3724). pCMV-HA-hIFITM plasmids were a gift from Jacob Yount and Howard Hang (Addgene plasmid # 58399; http://n2t.net/addgene:58399 ; RRID:Addgene_58399, Addgene plasmid # 58398; http://n2t.net/addgene:58398 ; RRID:Addgene_58398, Addgene plasmid # 58397; http://n2t.net/addgene:58397 ; RRID:Addgene_58397) (*66*).

### dsRNA immunoprecipitation

5×10^5^ VK2 cells in a 6-well plate were infected with HSV-2 at MOI 3 for 18 hours, lysed with 1 ml TBS (Thermo Fisher Scientific, J60877-K3) containing 0.5% NP-40 (Sigma-Aldrich, I8896-50ML), and centrifuged at 12,000g for 15 min. The cell lysates were incubated with 1 µg of mouse IgG control (Santa Cruz Biotechnology, sc-2025) or anti-dsRNA antibody (Cell Signaling Technology, 76651L) in the presence of 10 U of RNase inhibitor (Thermo Fisher Scientific, AM2694) at 4°C overnight. Dynabeads Protein G (Thermo Fisher Scientific, 10004D) was added to the mixtures, and incubated at 4°C for 2 hours. After four washes with the lysis buffer, the immunoprecipitates were isolated by RNA lysis buffer of High Pure RNA isolation kit (Roche, 11828665001). The RNA levels of the genes were analyzed by Brilliant III Ultra-Fast SYBR Green qRT-PCR Master Mix (Agilent Technologies, 600886). PCR primers: ND5 Forward, TCGAAACCGCAAACATATCA; ND5 Reverse, CAGGCGTTTAATGGGGTTTA; ND6 Forward, CCAATAGGATCCTCCCGAAT; ND6 Reverse, AGGTAGGATTGGTGCTGTGG; COX1 Forward, ACGTTGTAGCCCACTTCCAC; COX1 Reverse, TGGCGTAGGTTTGGTCTAGG; CYB Forward, AGACAGTCCCACCCTCACAC; CYB Reverse, GGTGATTCCTAGGGGGTTGT; ACTB, QuantiTect Primer Assay, QIAGEN, 249900, Hs01060665.

### MTT assay

VK2 human vaginal ECs were seeded in a 96-well plate at a density of 12×10^3^ cells/well. Cells were infected with HSV-2. For MTT assay, 10ul 98% Thiazolyl Blue Tetrazolium Bromide (Sigma) was added to each well and incubated for 4h. After 4h incubation, the medium was removed. Insoluble formazan was dissolved with DMSO. The plate was shaken for 10 minutes and read at 570nm wavelength. Percentage mitochondrial metabolic activity was calculated using the formula: 100 x ((A_sample_ – A_blank_) / (A_control_ – A_blank_)).

### Statistical analyses

For statistical analysis of data, we used 2-tailed Student’s t-test when the data exhibited normal distribution, and Wilcoxon rank-sum test when the data set did not pass the normal distribution test. When comparing more than 2 groups, multiple-comparison 1-way ANOVA was used with Tukey’s or Kruskal-Wallis multiple-comparison test or 2-way ANOVA with either Bonferroni’s or Sidak’s multiple-comparison test was used, as indicated in figure legends. For survival studies, the P values were calculated using log-rank (Mantel-Cox) test. For all experiments, P > 0.05 (NS); *0.01 < P < 0.05; **0.001 < P < 0.01; ***0.0001 < P < 0.001; ****P < 0.0001. All experimental data were reliably reproduced in 2 or more individual biological replicates. GraphPad Prism software or Stata was used for statistical analyses.

### Ethical approval

All animal experiments were approved in advance by the local authorities (Animal Ethics Committee at the Danish Veterinary and Food Administration (permission number 2021-15-0201-01084). All experiments were carried out in accordance with the Danish Animal Welfare Act for the Care and Use of Animals for Scientific Purposes, thus complying with local regulations. Human cervical samples were collected from women undergoing hysterectomy. All participants provided written informed consent, and the study was conducted under ethical approval granted by the Swedish Ethical Review Authority (Approval no. 2024-02834-01).

## SUPPLEMENTAL INFORMATION

Fig. S1 to S4 Table S1 and S2

## ACKNOWLEDGMENTS

Technical assistance of Ea Stoltze Andersen and is greatly appreciated. We also acknowledge Anja Bille Bohn from AU FACS Core facility, AU Health Bioimaging Core Facility, Mass Cytometry Unit (MCU), the Bioinformatics Core Facility, and animal core facility at Department of Biomedicine for the use of the facilities. The computational resources and support from GenomeDK and Aarhus University is acknowledged. The study was funded by the European Research Council (786602), the Lundbeck Foundation (R359-2020-2287), the Novo Nordisk Foundation (NNF23OC0084931), the Carlsberg Foundation (CF17-0687), the Danish National Research Foundation (CiViA, DNRF164), and the Swedish Research Council (2024-02549). Danish Agency for Higher Education and Science through the Danish national research infrastructure CellX (5229-00009B).

## AUTHOR CONTRIBUTIONS

L.S.R. and S.R.P. conceived the original project idea. X.L., L.S.R., and S.R.P. designed the study.

L.S.R. and S.R.P. led the development and management of the project. X.L., R.N., A.S., M.F.B., G.W., M.B.I., G.M., S.R.N.J., K.T., O.A.A., A.E. designed and/or performed experiments. K.H. and T.P.P.B, included the patients in the study and obtained patient cells and clinical information. X.D., F.D., and A.S. performed the bioinformatics analyses. X.L., X.D., R.N., A.S. F.D., C.B.V., K.B., L.S.R., and SRP analysed the data. S.R.P. acquired funding. S.R.P wrote the manuscript with inout from X.L. and L.S.R., and all authors reviewed and edited the paper.

### Data and material availability

All data needed to evaluate the conclusions in the paper are present in the paper or the Supplemental Materials. Further information and requests for resources and reagents should be directed to and will be fulfilled by the corresponding authors.

## Supplementary Figures

**Figure S1.**
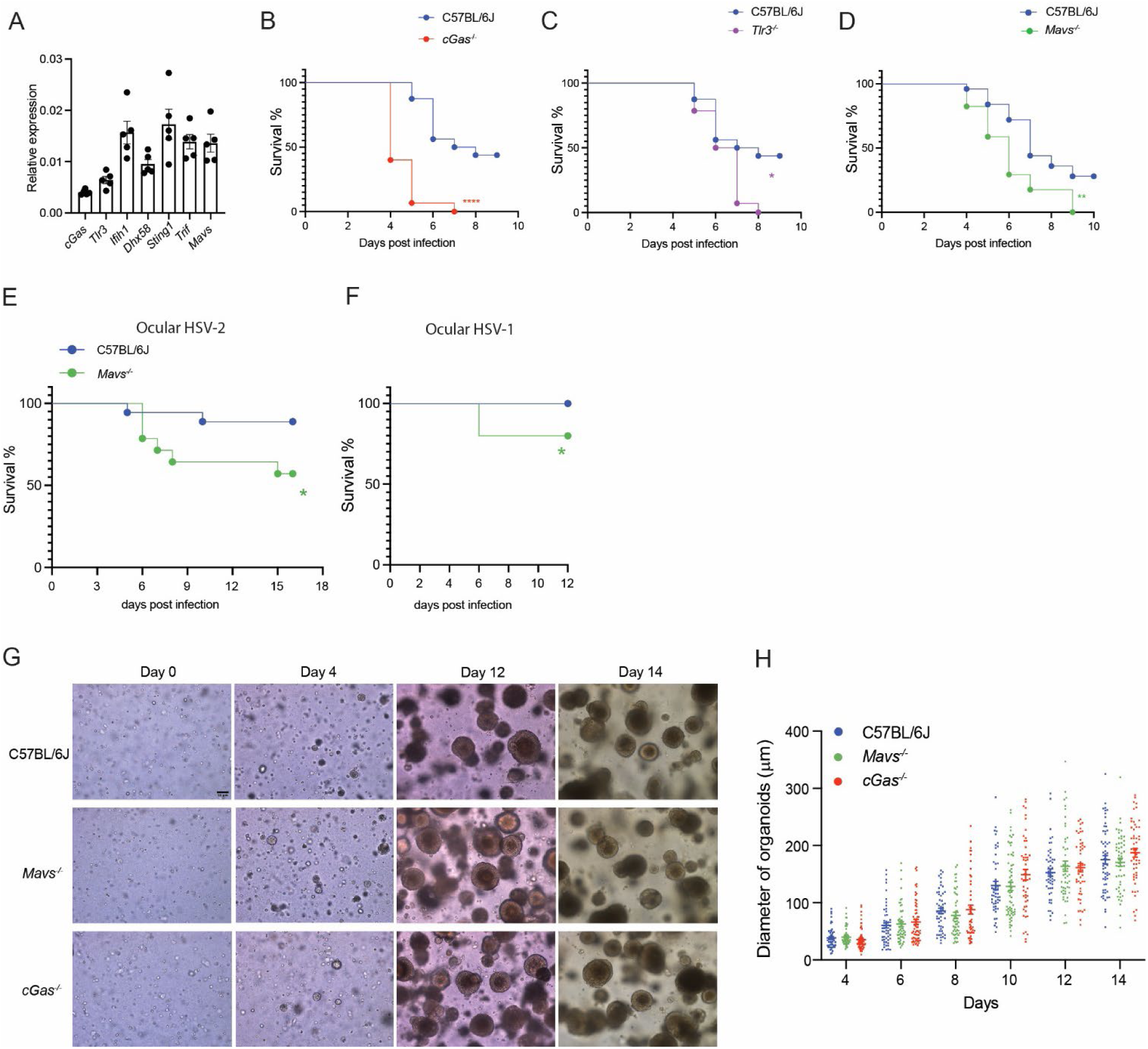
Host defense against genital HSV-2 relies on both RNA and DNA sensing pathways. (A) Relative expression in the mouse FGT of mRNAs encoding sensors and adaptors in the innate RNA and DNA sensing immune pathways. (B-D) Mice were infected intravaginally with 6.7 × 10^4^ plaque-forming units (p.f.u.) of HSV-2 (strain 333) and monitored on subsequent days for survival. (B) WT (C57BL/6) and *cGas^-/-^*(n = 8 mice/group), (C) WT and *Tlr3^-/-^* (n = 8 mice/group), and (D) WT and *Mavs^-/-^* (n = 6 mice/group). (E-F) WT and *Mavs^-/-^* mice were infected in the cornea with HSV-2 (200 p.f.u, n = 6-8 mice/group), or HSV-1 (2×10^6^ p.f.u, n = 7-8 mice/group) and monitored on subsequent days for survival. (G) Microscopy images of vaginal organoids from WT*, cGas^-/-^*, and *Mavs^-/-^* mice at the indicated time points seeding. (H) Measurement of the diameters of the vagina organoids from WT*, cGas^-/-^*, and *Mavs^-/-^* mice at the indicated time points post seeding. (B-F) Survival was analyzed using log-rank Mantel–Cox test.

**Figure S2.**
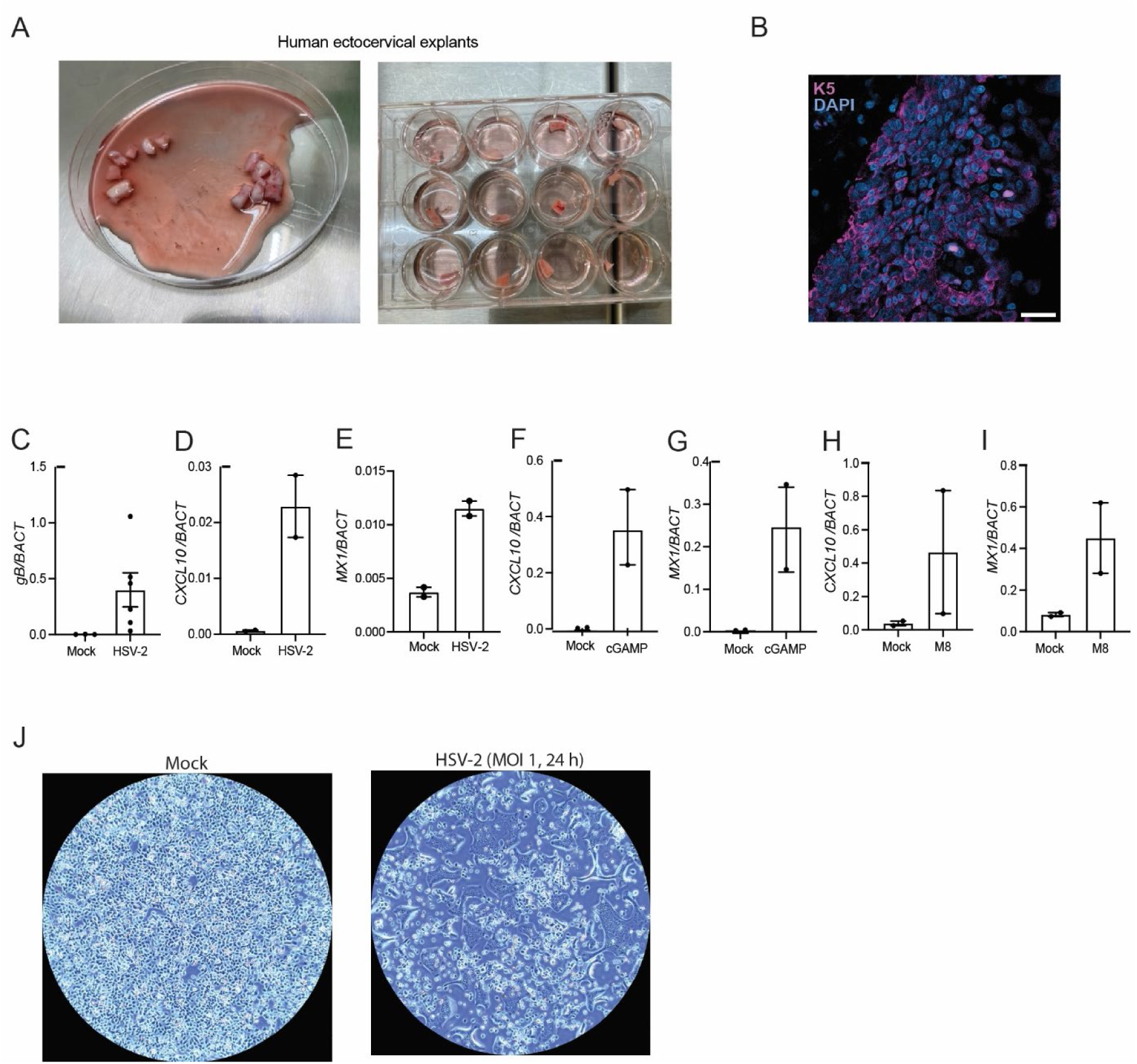
Human ectocervical explants. (A) Image of dissected ectocervical human tissue (left) and single tissue blocks in a cell culture 12-well plate. (B) Staining of sections from tissue blocks with anti-K5 (red) and DAPI (blue) and visualized with fluorescence microscopy. (C-I) mRNA levels of gB, *CXCL10*, or *Mx1* (normalized to housekeeping gene β-Actin) either after infection with HSV-2 (2×10^6^ p.f.u. 24 h), exposure to cGAMP (10 ug/ml, 24 h), or transfection with RIG-I agonist M8 (70 ng/ml, 24h). Each dot represents an explant. (J) Light microscopy images of VK2 cells treated with Mock or HSV-2 (MOI 1) for 24 h.

**Figure S3.**
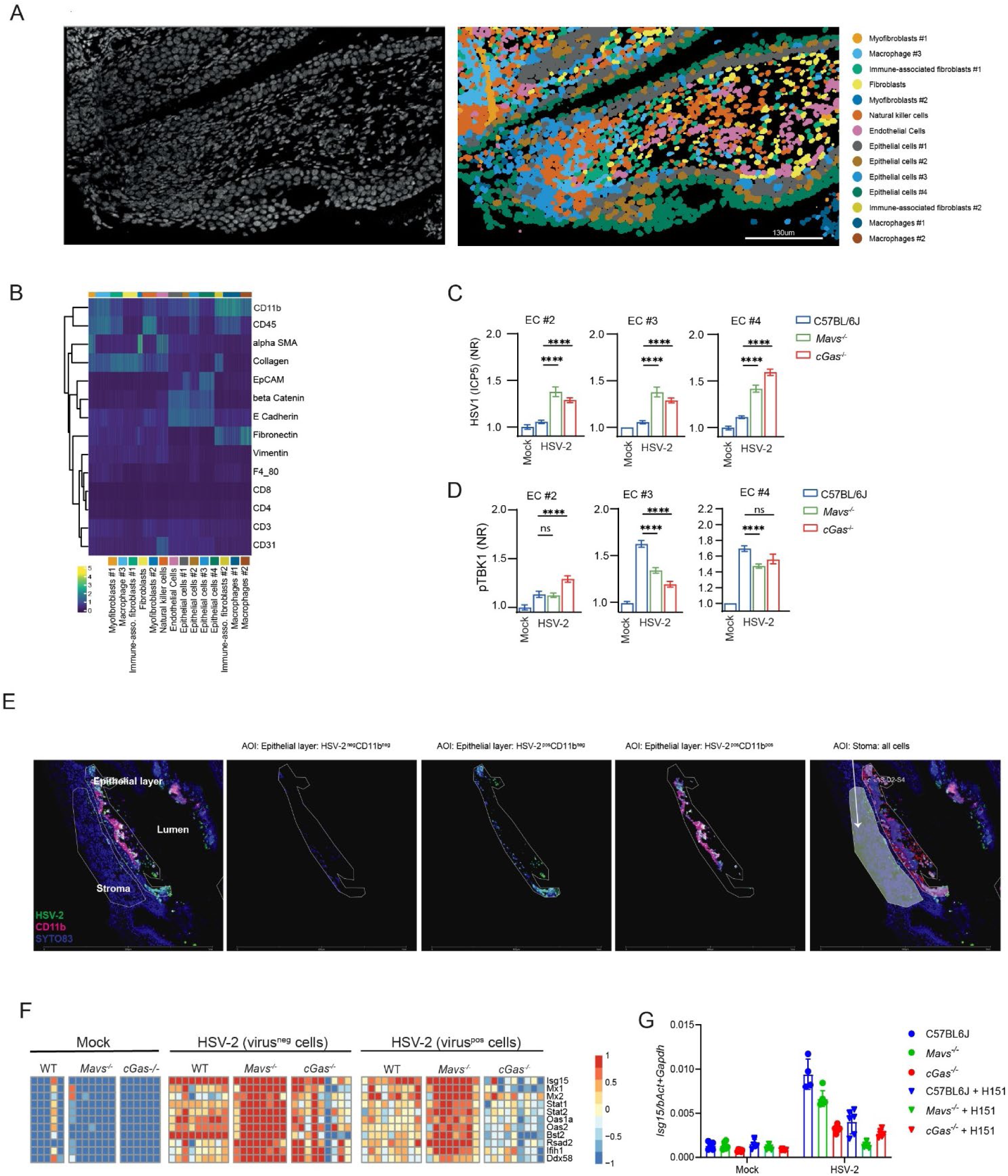
Response to genital HSV-2 infection in ECs. (A) Representative IMC image of DNA-Ir191/Ir193 DNA intercalator staining of vaginal tissue section from C57BL/6 mice (left), and overlaid cell segmentation with annotated cell types in defined colors (right). (B) Heatmap showing expression levels of markers used for annotation of cell types for IMC. (C) Quantification of HSV-2 (ICP5) in epithelial cell (EC) populations #2-4 in WT, *cGas^-/-^*, and *Mavs^-/-^* mice 48 h after infection (n=288-1980 cells/group, from 3-9 ROIs from IMC). (D) Quantification of pTBK1 in epithelial cell (EC) populations #2-4 in WT, *cGas^-/-^*, and *Mavs^-/-^* mice 24 h after infection (n=298-1254 cells/group, from 3-9 ROIs). (E) Representative image of Region of interest for transcriptomic analysis by GeoMx. (F) Heatmap of selected ISG transcripts in ECs analyzed with GeoMx digital spatial transcript profiler. HSV-2^+^ and HSV-2^-^ ECs were isolated from the vagina of mice infected for 48 h. (79 CD11b^-^ AOIs in the epithelial layers were analyze from 2-4 mice/group). (G) Levels of *Isg15* mRNA in vaginal organoids from WT*, cGas^-/-^*, and *Mavs^-/-^* mice 24 h after infection in the presence or absence of H151 (10 μM). Data were shown as means +/- st.dev. (C, D) The values are normalized to WT-mock group and *p* values were calculated using 2-way ANOVA with Bonferroni’s test. Plots show means ± st.dev; ^∗∗∗∗^*p* < 0.0001; ns, not significant. All mice in this figure were infected intravaginally with HSV-2 (6.7 × 10^4^ p.f.u. per mouse).

**Figure S4.**
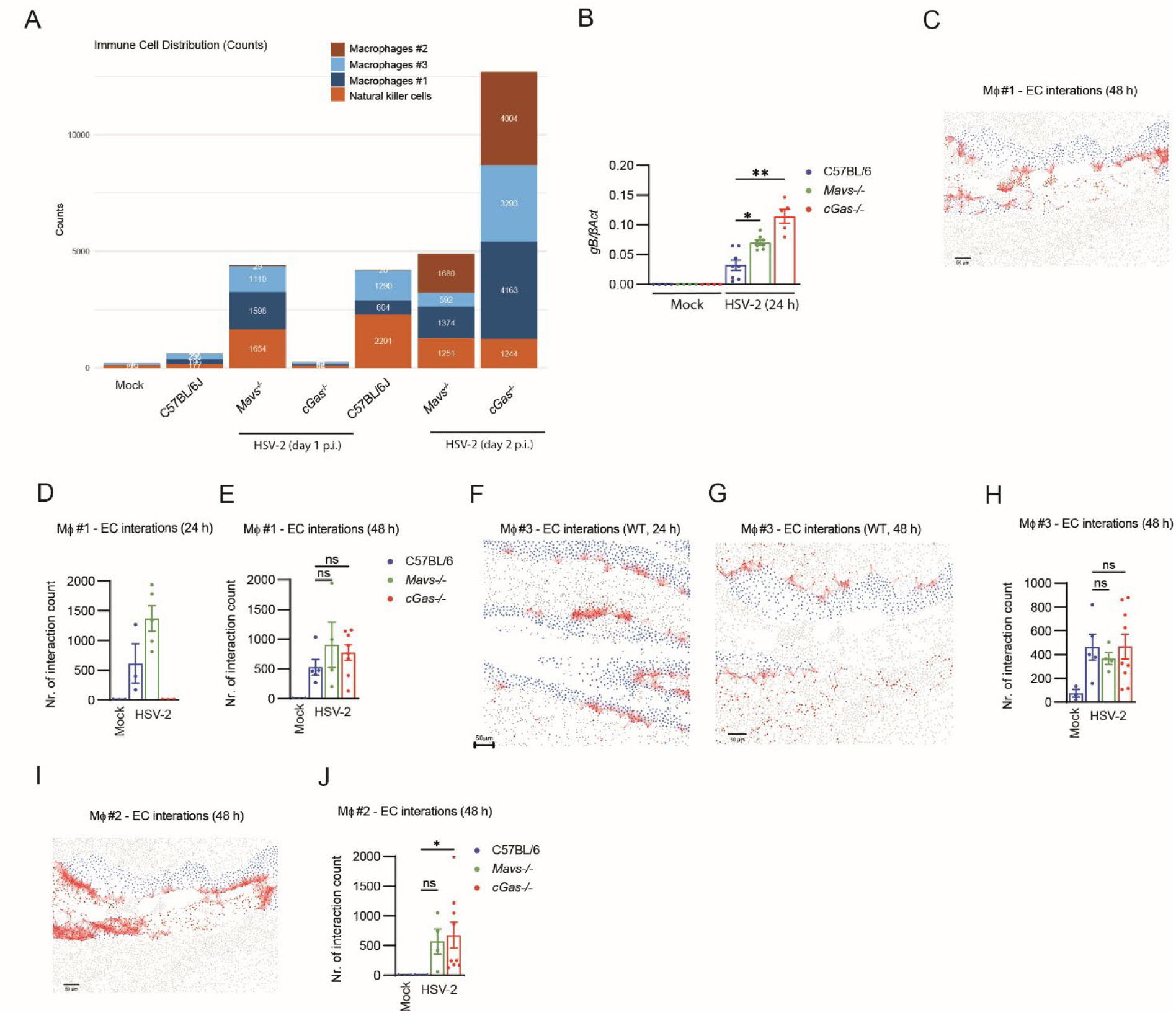
Recruitment and activation of leukocytes in response to HSV-2 infection in the FGT. (A) Quantification by IMC of number of leukocytes in the annotated immune cell populations at the indicated time points after vaginal infection of WT, *cGas^-/-^*, and *Mavs^-/-^* mice. (B) HSV-2 gB transcript levels in total vagina RNA isolated 24 h infection from WT, *cGas^-/-^*, and *Mavs^-/-^* mice. (C) Representative IMC image of spatial interaction analysis for macrophage #1 and ECs in the vagina of WT mice 48 h after vaginal infection. Spatial interaction analysis based on proximity (≤ 30 µm), Mφ#1 (red), EC (blue), and other cells (gray). Scalebar, 50 μm. (D-E) Quantification by IMC of macrophage #1 - EC interactions in the vagina of WT, *cGas^-/-^*, and *Mavs^-/-^*mice 24 and 48 h after HSV-2 infection. Each dot represents one ROI. (F-G) Representative images of spatial interaction analysis for macrophage #3 and ECs in the vagina of WT mice 24 and 48 h after vaginal infection. Spatial interaction analysis based on proximity (≤ 30 µm), Mφ#3 (red), EC (blue), and other cells (gray). Scale-bar, 50 mm. (H) Quantification of macrophage #3 - EC interactions in the vagina of WT, *cGas^-/-^*, and *Mavs^-/-^*mice 48 h after HSV-2 infection. Each dot represents one ROI. (I) Representative image of spatial interaction analysis for macrophage #2 and ECs in the vagina of *cGas^-/-^* mice 48 h after vaginal infection. Spatial interaction analysis based on proximity (≤ 30 µm), Mφ#2 (red), EC (blue), and other cells (gray). Scalebar, 50 μm. (J) Quantification of macrophage #2 - EC interactions in the vagina of WT, *cGas^-/-^*, and *Mavs^-/-^* mice 48 h after HSV-2 infection. Each dot represents one ROI. (B, E, H, J) *P* values were calculated using ANOVA with a Turkey’s post hoc test. ^∗^*p* < 0.05; ^∗∗∗^*p* < 0.01; ns, not significant. All mice in this figure were infected intravaginally with HSV-2 (6.7 × 10^4^ p.f.u. per mouse).

